# An MRI-informed histo-molecular analysis implicates ependymal cells in the pathogenesis of periventricular pathology in multiple sclerosis

**DOI:** 10.1101/2025.01.14.633055

**Authors:** Adam M.R. Groh, Elia Afanasiev, Risavarshni Thevakumaran, Liam Callahan-Martin, Finn Creeggan, Moein Yaqubi, Stephanie Zandee, Alexandre Prat, David A. Rudko, Jo Anne Stratton

## Abstract

It is now widely recognized that the cerebrospinal fluid (CSF)-adjacent brain surfaces – namely the subpial cortical region and the ependyma-adjacent periventricular region – are uniquely susceptible to a distinct, diffuse form of pathology in multiple sclerosis. So-called *surface-in* gradients of pathology predict future disease relapses independent of classical white matter lesions and are thought to occur as a result of cytotoxic factors in the CSF. Given the underlying mechanisms driving *surface-in* gradients appear to be distinct, they represent a novel treatment target. However, exactly how cytotoxic factor entry into the brain is regulated at these CSF-facing borders is not understood, particularly at the ventricular interface. Indeed, although studies have indicated that ependymal cells may be damaged in MS, there has yet to be a comprehensive assessment of cell health in the disease. We employed ultra-high-field MRI-guided immunohistochemistry, electron microscopy, and multiomic single nucleus RNA/ATAC sequencing to deeply phenotype human ependymal cells in MS. Our data revealed that ependymal cell pathology is a direct correlate of periventricular *surface-in* gradients of pathology in MS, and that the immune-responsive, reactive state assumed by ependymal cells is associated with widespread transporter and junctional protein gene dysregulation. We then further defined the gene regulatory networks underpinning the MS ependymal state, predicted ligands known to be enriched in MS CSF that could drive the emergence of this state, and tested one candidate *in vivo*. We found that IFNγ increased murine ependymal permeability and that conditional knockout of ependymal interferon gamma receptor 1 (Ifngr1) was sufficient to reverse this effect. Our data directly implicate ependymal cell dysregulation in the emergence of periventricular pathology in MS. More widely, we denote the modulatory capacity of CSF ligands on ependymal cell function and how this may influence the inflammatory status of the periventricular region.

**Graphical Abstract:** 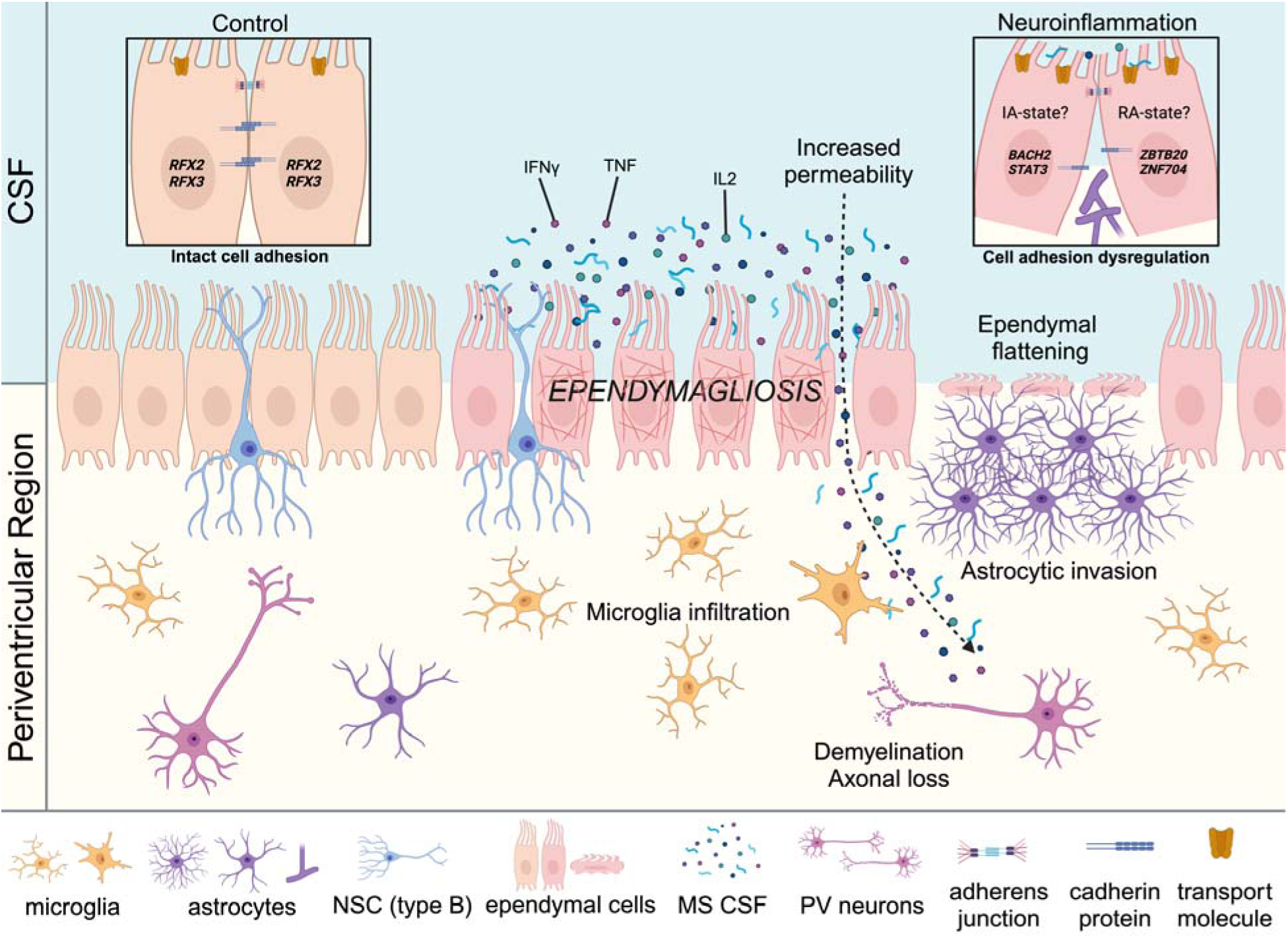

## Introduction

Multiple sclerosis (MS) is an autoimmune disease characterized by central nervous system (CNS) inflammation, demyelination, and neurodegeneration.^1,2^ MS pathology has classically focused on focal lesions which form as a result of blood brain barrier (BBB) breakdown and immune cell entry.^3^ Lesions typically associate with post-capillary venules and can be categorized based on the pattern and extent of demyelination and immune cell infiltration.^3^ Although studies of focal pathology have dominated the literature, diffuse pathology is also pervasive throughout white and grey matter structures of the MS brain^4–7^ and can be challenging to identify by standard *in vivo* MRI. ^5,6^ So-called *surface-in* gradients of pathology exist adjacent to cerebrospinal fluid (CSF)-exposed brain areas, such as the subpial and periventricular regions, and dissipate with distance from the CSF.^6,8^ Indeed, thalamic damage observed in MS is most pronounced at the surface of the ventricles,^9,10^ and periventricular deep grey matter damage also correlates with meningeal CSF inflammation and damage to other CSF-facing grey matter regions.^4,11^ Most importantly, however, is that *surface-in* gradients of pathology directly correlate with disability^12–14^ and can independently predict disease relapses.^15,16^

The relationship of *surface-in* gradients to CSF-facing brain borders is highly suggestive of cerebrospinal fluid (CSF) involvement in disease pathogenesis.^6,8,9^ Several histopathological studies have demonstrated that the leptomeninges of MS brain tissue are populated by aggregates of stromal and immune cells, which form ectopic follicles known as tertiary lymphoid tissues (TLTs).^8,17–19^ It has been proposed that meningeal TLTs release pro-inflammatory molecules into the CSF that diffuse across the CSF-parenchyma interface where they can cause demyelination and brain atrophy.^4,20^ This hypothesis is particularly compelling given the CSF of MS patients is well-described to have elevated levels of immune cells, proinflammatory cytokines, complement, and toxic coagulation factors like fibrinogen.^8,11,21–25^

While proinflammatory molecules in the CSF provide a reasonable explanation for the emergence of *surface-in* gradients, the specific mechanism(s) by which cytotoxic factors gain access to the parenchyma remain less clear. Within the periventricular region, ependymal cells form a specialized interface separating the CSF from the CNS parenchyma,^26^ and evidence suggests that they may be damaged in MS^27,28^ and in preclinical models of the disease.^29^ Despite this, there has been no comprehensive examination of the ependyma with respect to neighboring periventricular gradient pathology in MS, and no molecular dissection of the mechanisms that could be driving this damage.

Here, we interrogated the relationship between human ependymal cell pathology and periventricular pathology by applying ultra-high-field (7T) *ex vivo* MRI-guided immunohistochemistry. We identified periventricular *surface-in* gradients in white matter of MS brains and confirmed histologically that these regions showed demyelination, axonal loss, and increased microglial presence. Importantly, *ependymagliosis* (ependymal cellular reactivity) was largely restricted to these sites. To further understand the functional impact of ependymal reactivity, we conducted multiomic single nucleus RNA/ATAC sequencing on fresh-frozen human periventricular brain tissue and noted considerable dysregulation of transporter activity and junctional protein-associated genes in MS ependymal cells. Ultrastructural analysis confirmed that ependymal cell-cell adhesion was altered in periventricular lesions in MS. We then defined the gene regulatory networks (GRNs) underpinning the transcriptomic shift of normotypic ependymal cells to an MS-associated state and predicted ligands capable of driving the emergence of this state. IFNγ – a top candidate ligand – was found to modulate ependymal barrier capacity *in vivo*, and conditional knockout of the ependymal interferon gamma receptor 1 (Ifngr1) reversed this effect.

Altogether, we provide evidence of a phenomenon in which proinflammatory factors in the CSF modulate ventricular wall permeability via direct alterations to ependymal cell-cell adhesion, which may impact the inflammatory status of the ventricular-subventricular zone (VZ-SVZ) in MS and other neuroinflammatory disorders of the CNS that have altered CSF composition.

## Results

### Ultra-high-field ex-vivo MRI reveals periventricular surface-in gradients in white matter of multiple sclerosis brains

Recent MRI studies have uncovered non uniform abnormalities in white and grey matter that are most pronounced directly adjacent to the glia limitans superficialis at the pia surface and the ependyma at the ventricular surface.^6^ These abnormalities decrease with distance from the CSF^5,6,15,30,31^ — so called *surface-in* gradients of pathology. To date these studies have primarily been conducted *in vivo* and/or are limited by time and field strength, making the possibility of detecting all abnormalities, however subtle, a challenge. Here we exposed ex-vivo formalin-fixed human MS brain slabs (n=12) and age- and sex-matched non-inflammatory, non-neurodegenerative disease control brain slabs (n=11) (Supp. Table 1) to ultra-high-field (7T) MRI to identify areas of periventricular pathology. Once scanned, each brain image (Figure 1Ai) was subdivided into angular bands (15° angular coverage) (Figure 1Aii) and concentric, geodesic bands (1.5mm wide) (Figure 1Aiii). Our tissue segmentation methodology permitted a rigorous and unbiased quantitative approach for incremental analysis of T1 and T2* relaxation time values (averaged along the entire depth of each slab) from the ventricular surface to the cortical surface (ependyma-in) within each angular band, which was necessary for subsequent systematic selection of tissue blocks (Figure 1Aiv) for histological analysis (Figure 1A). Representative plots of T1 and T2* relaxation time values in four angular bands in a single ex-vivo control brain slab illustrate the uniform relaxation time response across all geodesic bands, indicative of normal periventricular myelin (Figure 1Bi). Conversely, plots of T1 and T2* relaxation time in four angular bands from a representative ex-vivo MS brain slab demonstrate prominent increases in relaxation time adjacent to the ventricle (geodesic band 2) that gradually decreased to a level comparable to control by geodesic band 10 (Figure Bii). In 11 out of 12 MS brain slabs and 9 out of 10 control slabs, there were angular bands exhibiting a gradient of reduction in relaxation time from the ventricular surface outwards (ependyma-in), although the size of these gradients varied considerably between control and MS slabs. We therefore assessed the total change in periventricular white matter T1 and T2* across geodesic bands (ependyma-in) by determining angular bands exhibiting an ependyma-in reduction in T1 or T2*. In this way, we were able to plot the exact values of all T1 and T2* ependyma-in reductions in n=61 MS angular bands and n=59 control angular bands (Figure 1C; Supp. Table 2). These data revealed a significant increase in the total change, with respect to the ventricular surface, in T1 (Mann-Whitney, p = 5.0e-10) and T2* (Mann-Whitney, p = 5.2e-06) in MS compared to control (Figure 1C). Through this approach, we identified angular/geodesic bands with pronounced or severe gradient periventricular myelin abnormalities in MS that were extracted for further histological analysis. We also stratified this data by slab ID to ensure this was a consistent finding across MS slabs and not driven by a particular brain slab or patient (Figure 1C). Interestingly, there were rare instances of equally severe periventricular gradient abnormalities in control brains (across two separate donors), which can occur with aging,^10,32^ but we were unpowered to characterize the relationship of these abnormalities with histological analyses.

**Figure 1.**
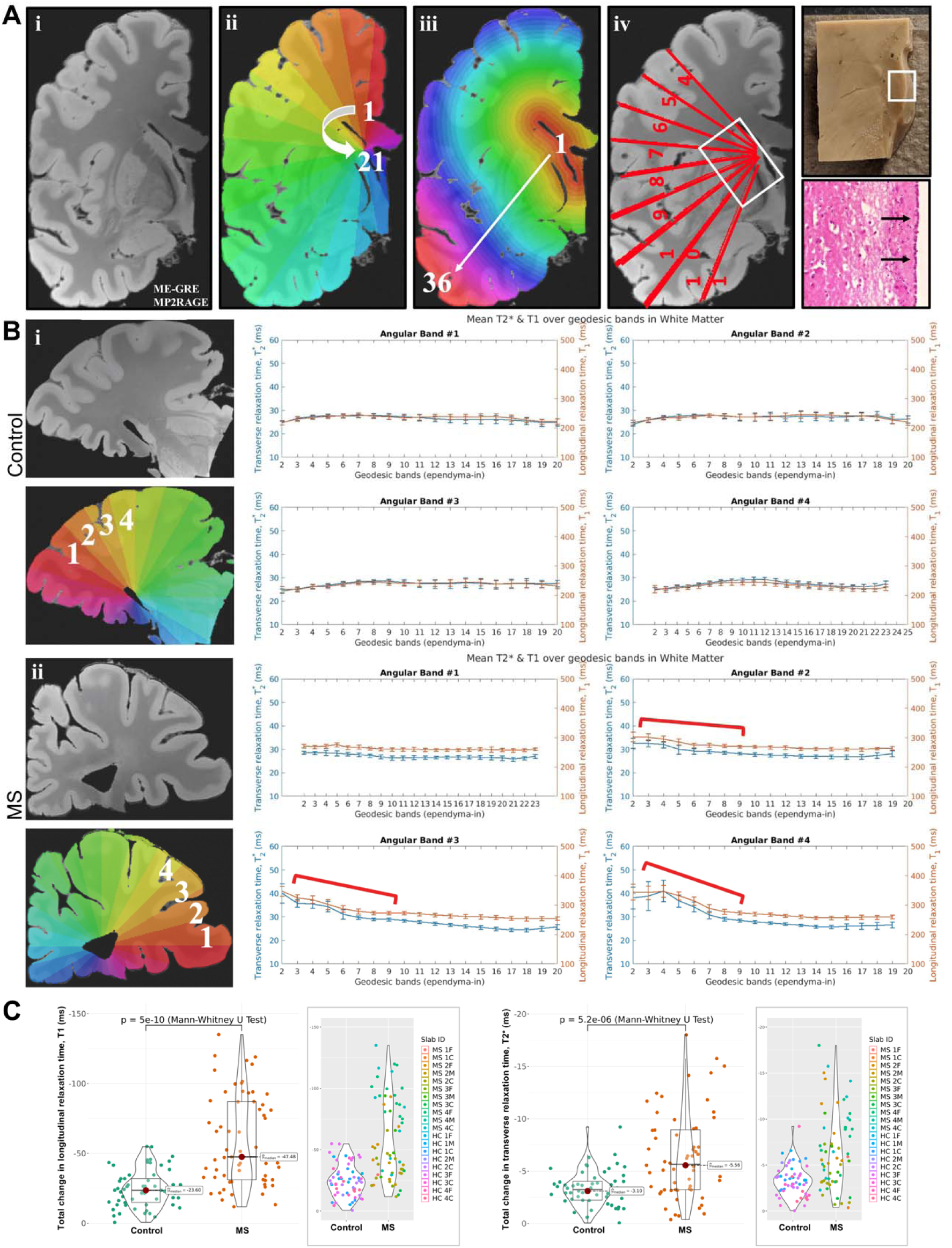
Ultra-high-field (7T) MRI reveals the presence of surface-in periventricular gradients in white matter of *ex vivo* MS brain tissue. **(A)** Schematic of the ultra-high-field MRI-guided immunohistochemistry workflow employed in the study. **(i)** Representative 2D image of a scanned *ex vivo* formalin-fixed human brain slab. 3D ME-GRE (T2*) and MP2RAGE (T1) sequences were employed to assess myelin integrity. **(ii)** Representative image of brain slab scan subdivision into 21 angular bands (15° angular coverage); the number of angular bands per brain slab varied with brain slab size. **(iii)** Representative image of brain slab subdivision into 36 concentric, geodesic bands (1.5mm wide); the number of geodesic bands per brain slab varied with brain slab size. **(iv)** Representative image of periventricular tissue block selection methodology. Periventricular tissue blocks containing specific angular bands (here, bands 4-11) in which there was evidence of gradients in T2* and T1 (transverse and longitudinal relaxation time) across geodesic bands were resected, in addition to blocks that did not contain evidence of gradients in T2* and T1. This permitted downstream histological evaluation of both gradient-negative (GR-) and gradient-positive (GR+) periventricular regions in control and MS human brain slabs. **(B) (i)** Representative 2D image, and associated transverse and longitudinal relaxation time plots, of a control *ex vivo* formalin-fixed human brain slab. In this slab, angular bands 1-4 showed no evidence of gradients in transverse or longitudinal relaxation time from the periventricular surface, outwards (geodesic bands 2-20). **(B) (ii)** Representative 2D image, and associated transverse and longitudinal relaxation time plots, of an MS *ex vivo* formalin-fixed human brain slab. In this slab, angular bands 2-4 showed evidence of gradients in transverse and longitudinal relaxation time (marked by red brackets) from the periventricular surface, outwards (geodesic bands ∼2-20). **(C)** Violin plots denoting the total change in longitudinal relaxation time (T1; ms; left) and transverse relaxation time (T2*; ms; right) in all resected blocks (GR- and GR+) from control and MS brains. There was a significant increase in total change in both longitudinal (p=5e-10) and transverse relaxation time (p=5.2e-06) in MS blocks compared to control blocks, demonstrating an increased load of periventricular (*surface-in*) pathology. On the right of each violin plot is a second violin plot with the data stratified by patient, denoting that all patients showed this phenotype.

### Periventricular white matter surface-in gradients show evidence of demyelination, increased microglial density, axonal loss, and are lined by reactive ependymal cells

It is well documented that subpial gradients of neuronal loss and demyelination occur in the cortical grey matter in MS and correlate with the presence of meningeal inflammation.^4^ Similar gradients are also present in the deep grey matter (thalamus) of MS brains with pathology being more severe adjacent to the ependyma.^10^ Although evaluated by MRI, periventricular white matter regions have not been histologically assessed in this context, nor has ependymal pathology. To systemically select tissue segments for histological analysis, we used two criteria. Firstly, we selected segments if the distributions of T1 and T2* relaxation times at the first and last geodesic band, used to compute total change in relaxation time (Supp. Table 2), were found to be significantly different by Wilcoxon rank-sum test (p<0.05). Secondly, we selected segments if the absolute value of the total reduction in T1 and T2* was greater than threshold values of |ΔT1|>30.0ms and |ΔT2*|>2.0ms respectively. The threshold values of |ΔT1| and |ΔT2*| were determined, in this context, based on inspection of all ependyma-in reductions reported in Supp. Table 2. Using this approach, we resected tissue blocks containing evidence of severe gradient abnormalities (GR+) and regions without pronounced gradient abnormalities (GR-) in control (n = 2 GR+; n = 16 GR-) and MS (n = 16 GR+; n = 12 GR-) brains. These MRI-identified tissue blocks were then stained to investigate the nature of periventricular pathology and ependymal dysregulation. Hematoxylin and Eosin (H&E) staining was used to confirm the presence and location of ependymal cells in each tissue block (Figure 1A). Luxol fast blue analysis (Figure 2A) reaffirmed T1 and T2* relaxation time data, demonstrating a significant loss of periventricular myelin in close proximity to the CSF (1mm) as measured by color intensity in MS GR+ blocks compared to GR- control (p<0.01) and GR- MS (p<0.05) blocks (Figure 2B). Again, consistent with our MRI data, periventricular demyelination was present near the CSF (1mm) in GR+ MS blocks, which subsided distally (3mm) (Figure 2B); whereas there was no change in myelin between 1mm and 3mm in the GR- control blocks (Figure 2B). Microglia density was increased in both GR- and GR+ MS blocks compared to GR- control blocks (p>0.05; Figure 2C-D). There was also a significant increase in microglia density near the CSF (1mm) in GR+ and GR- MS blocks which diminished distally (3mm) (p<0.01), and this gradient effect was not seen in GR- control brain blocks (p=0.74) (Figure 2D). Importantly, there was no difference in the density of microglia between the MS GR- and MS GR+ blocks (Figure 2D). As for axonal analysis, GR+ MS blocks had a significant decrease in axonal density compared to GR- control (p<0.01) and GR- MS blocks (p<0.05) (Figure 2E-F). Loss of axons resolved distally (3mm) from the CSF in GR+ MS blocks (P<0.05), whereas there was no gradient effect in GR- control blocks (p=0.70) (Figure 2F).

**Figure 2.**
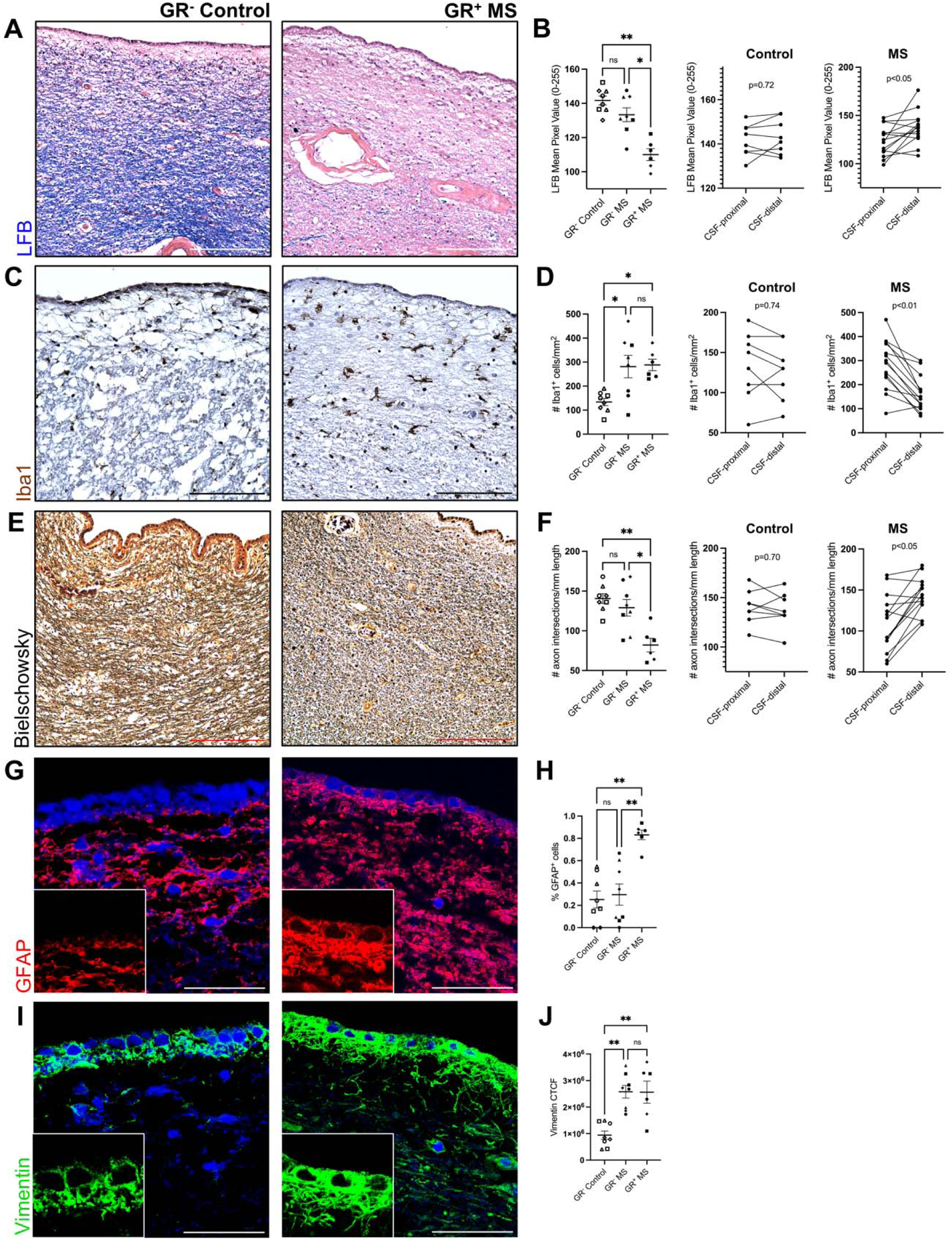
MRI-guided immunohistochemistry confirms that gradient-containing regions show evidence of demyelination, increased presence of microglia, axonal loss, and are lined by reactive ependymal cells. **(A)** Representative luxol fast blue (LFB) images showing decreased staining intensity in GR+ blocks in MS compared to GR- blocks in control. Scale bars = 200µM. **(B)** Quantification of LFB mean pixel value in GR- blocks in control compared to GR- and GR+ blocks in MS, proximal to the CSF (1mm) (left side), showing decreased LFB mean pixel values in GR+ regions in MS compared to GR- regions in MS (p<0.05) and control (p<0.01). LFB mean pixel values in CSF-proximal regions (1mm) were also compared to a CSF-distal regions (3mm); these data are plotted separately for control and MS blocks (right side). This analysis demonstrated no change in LFB mean pixel value between CSF-proximal and CSF-distal regions in control (p=0.72), whereas there was an increase in LFB mean pixel value in CSF-distal regions compared to CSF-proximal regions in MS (p<0.05), suggesting increased myelination. **(C)** Representative Iba1 images showing increased microglia presence in GR+ blocks in MS compared to GR- blocks in control. Scale bars = 200µM. **(D)** Quantification of the number of Iba1+ cells/mm^2^ in GR- blocks in control compared to GR- and GR+ blocks in MS, proximal to the CSF (1mm) (left side), showing increased numbers in GR+ and GR- blocks in MS compared GR- blocks in control (p<0.05). The number of Iba1+ cells/mm^2^ in CSF-proximal regions (1mm) was also compared to CSF-distal regions (3mm); these data are plotted separately for control and MS blocks (right side). This analysis demonstrated no change in the number of Iba1+ cells/mm^2^ between CSF-proximal and CSF-distal regions in control (p=0.74), whereas there was a decrease in the number of Iba1+ cells/mm2 in CSF-distal regions compared to CSF-proximal regions in MS (p<0.01), suggesting less microglial infiltration. **(E)** Representative Bielschowsky images showing fewer axonal intersection/mm length in GR+ blocks in MS compared to GR- blocks in control. Scale bars = 200µM. **(F)** Quantification of the number of axons/mm length in GR- blocks in control compared to GR- and GR+ blocks in MS, proximal to the CSF (1mm) (left side), showing decreased numbers in GR+ blocks in MS compared GR-blocks in MS (p<0.05) and control (p<0.01). The number of axons/mm length in CSF-proximal regions (1mm) was also compared to CSF-distal regions (3mm); these data are plotted separately for control and MS blocks (right side). This analysis demonstrated no change in the number of axons/mm length between CSF-proximal and CSF-distal regions in control (p=0.70), whereas there was an increase in the number of axons/mm length in CSF-distal regions compared to CSF-proximal regions in MS (p<0.05), suggesting increases numbers of axons. **(G)** Representative GFAP images showing an increased percentage of GFAP+ ependymal cells (*ependymagliosis*) lining GR+ blocks in MS compared to GR- blocks in control. Scale bars = 40µM. **(H)** Quantification of the percentage of GFAP+ ependymal cells lining GR- blocks in control compared to GR- and GR+ blocks in MS, showing an increase in the percentage of GFAP+ ependymal cells lining GR+ blocks in MS compared to GR- blocks in MS (p<0.01) and control (p<0.01). **(I)** Representative Vimentin images showing increased expression by ependymal cells (*ependymagliosis*) lining GR+ blocks in MS compared to GR- blocks in control. Scale bars = 40µM. **(J)** Quantification of ependymal Vimentin expression intensity lining GR- blocks in control compared to GR- and GR+ blocks in MS, showing an increase in Vimentin corrected total cell fluorescence (CTCF) in ependymal cells lining GR+ and GR- blocks in MS compared to GR- blocks in control (p<0.01). *p<0.05; **p<0.01. Unfilled squares, triangles, circles, and diamonds represent individual control patients (H1, 2, 3 and 4, respectively; Supp. Tables 1-2). Filled triangles, squares, circles, and diamonds represent individual MS patients (MS1, 2, 3, and 4, respectively; Supp. Tables 1-2).

Multiciliated mature ependymal cells do not typically express glial fibrillary acidic protein (GFAP). However, akin to astrocytes, they can undergo a process we termed *ependymagliosis* when stressed,^33^ upregulating proteins associated with classical descriptions of astrogliosis. This includes GFAP^34^ and vimentin, two cytoskeletal proteins known to be upregulated in ependymal cells in response to various brain injuries.^34–36^ To evaluate the ependyma, we stained periventricular tissue blocks with GFAP (Figure 2G-H) and vimentin (Figure 2I-J). There were significantly higher proportions of GFAP-positive ependymal cells lining the ventricles of GR+ MS blocks compared to GR- control (p<0.01) and GR- MS blocks (p<0.01) (Figure 2H). Ependymal vimentin fluorescence intensity was also increased, but in both GR+ and GR- MS blocks compared to GR- control blocks (p<0.01) (**Figure 2J**). This suggests that although some features of *ependymagliosis* (i.e., increased vimentin expression) may be initiated in regions where there is no *surface-in* gradient pathology in MS (GR- MS), it has not reached a state of widespread cytoskeletal protein dysregulation (where both GFAP and vimentin are affected, as in GR+ MS regions). These data demonstrate that pronounced *ependymagliosis* is highly specific to regions of *surface-in* gradients of demyelination and axonal loss in MS, but that subtle *ependymagliosis* is already underway in gradient-free regions.

### Single nucleus RNA sequencing of human ependymal cells in MS confirms the existence of a reactive transcriptomic state associated with cell adhesion protein gene dysregulation

Having provided correlative evidence that sites of *surface-in* gradients of periventricular pathology are lined by reactive ependymal cells (suggestive of their involvement of the emergence of this pathology) we wanted to more precisely investigate the nature of ependymal cell dysregulation through a comprehensive assessment of their transcriptome and epigenome. To do this, we extracted samples of periventricular white matter from archived frozen human brain tissue sourced from individuals with MS (n=5) and age and sex-matched controls (n=4) for multiomic single nucleus RNA/ATAC sequencing (Figure 3A). In order to increase the likelihood that we resected ependymal cells from regions adjacent to both normotypic and pathological periventricular parenchyma, we systematically dissected tissue from both the medial and lateral walls of the lateral ventricle from one rostral and one caudal region (Figure 3A). Importantly, 4/5 patients used for sequencing were previously shown to have periventricular pathology in the ipsilateral hemisphere as per Figures 1 and 2. We captured a variety of periventricular cell types, including ependymal cells, and confirmed that most cell types were present in each patient we sequenced (Figure 3B). We confirmed brain cell identities by evaluating the expression of cell type-specific genes, which was *FOXJ1* for ependymal cells (Figure 3C). Ependymal cells were subset from the integrated periventricular object for all downstream analyses. We employed a machine-learning informed package termed RePACT^37^ to predict genes with the most variable expression between control and MS-associated ependymal cells. We then generated a 3D PCA plot depicting a clear transcriptomic shift of ependymal cells in a control state (blue) towards an MS-associated state (red) (Figure 3D). Genes associated with the control ependymal cell state were involved in ciliary regulation (*PIFO*), calcium binding (*CALM1*), mitochondrial respiration and energy homeostasis (*HIGD2A*, *ATP5MG, CKB*), solute transport (*SLC14A1*), and mRNA and protein transport (*NPIPA1*, *MYL6*) (Figure 3E; Supp. Figure 1A). Genes associated with the MS ependymal cell state were involved in cell-cell recognition and adhesion (*PCDH7*), interferon responsivity (*IFI16*), regulation of cell size, shape, and proliferation (*PRR16, SIPAL1, PTPN9*), solute transport (*SLC1A2*), and synaptic regulation and neuronal guidance (*ROBO1*, *EPHA5*) (Figure 3E; Supp. Figure 1A). We then performed gene ontology analysis using the variable gene output from RePACT, and confirmed similar results as predicted by classic differential gene expression-informed gene ontology (Figure 3F; Supp. Figure 1B). Indeed, MS ependymal cells upregulated a variety of genes associated with cell adhesion and transporters (Figure 3F), suggesting that the machinery necessary for maintaining fluid equilibrium (cell-cell adhesion and transporter proteins) between the CSF and the periventricular parenchyma may be altered in MS. Further, a variety of GO terms associated with synaptic organization and specification were enriched in MS ependymal cells (Figure 3F). We suggest that upregulation of synapse-associated genes is an artifactual association, in the sense that ependymal cells likely express synapse-associated adhesion proteins for alternative purposes, such as to control cell-cell adhesion. Alternatively, this genetic regulation could indicate changes in the interaction of ependymal cells with supraependymal serotonergic axons (which have been predicted, yet not conclusively proven, to alter multiciliated ependymal cell functions under homeostatic conditions).^38^ When we profiled individual differentially expressed genes further, we found that a gamut of adhesion protein-associated genes (i.e., catenin family, cadherin family, and other adhesion-associated glycoproteins) were significantly upregulated (p.adj < 0.05) in MS ependymal cells compared to control (Figure 3G), which we interpreted as compensatory genetic regulation to substantiate cell-cell contact. We also saw clear upregulation of some transport molecules like *AQP4* and *SLC1A2* (Figure 3G). Given the direct potential relevance of altered ependymal cell adhesion to the emergence of the CSF-mediated *surface-in* gradients of periventricular pathology we observed previously (Figures 1 and 2), we focused on phenotyping this phenomenon further.

**Figure 3.**
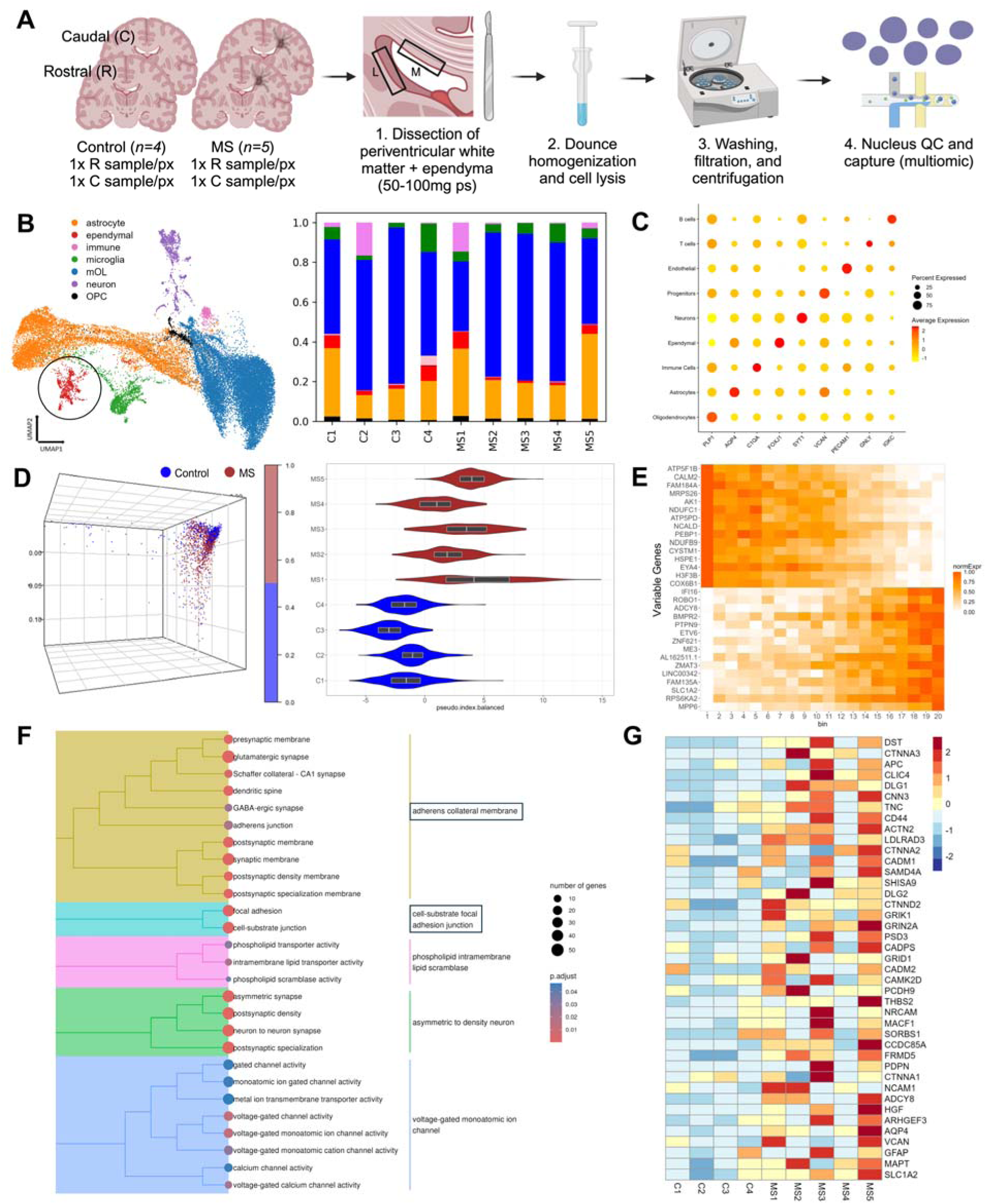
Single nucleus RNA sequencing of human ependymal cells in MS confirms the existence of a reactive transcriptomic state associated with cell adhesion protein gene dysregulation. **(A)** Schematic representation of tissue dissection and multiomic sequencing workflow. R = rostral; C = caudal; px = patient. **(B)** (Left) UMAP visualization of cell types identified through multiomic sequencing. Ependymal cells (red) are denoted by a black circle and were subset for further analysis. (Right) Stacked bar plot showing normalized proportions of cell types per donor. Ependymal cells were captured in all donors per experimental group (control and MS). **(C)** Dot plot of periventricular cell type-associated gene expression. Dot colour indicates the average expression value, whereas dot size indicates the percentage of cells expressing the gene. **(D)** (Left) 3D PCA plot demonstrating the control (blue) to MS (red) ependymal cell trajectory following a logistic regression-PCA-derived analysis. (Right) Violin plot showing a representation of overall disease score per patient, following a logistic regression-PCA-derived analysis. All MS patients showed an increase in pseudo-disease index compared to control patients, although there was clear variability within groups. **(E)** Heatmap showing normalized expression of the top 15 statistically significant genes (q-value < 0.05) per condition (control – top 15; MS – bottom 15) along a logistic regression-PCA-derived MS trajectory, organized into low-to-high binned pseudostates. Bin 1 represents the least diseased cells (control), while bin 20 represents the most diseased cells (MS). Note an increase in the expression of genes associated with response to interferons and cell adhesion in the most diseased (bin 20) cells (*IFI16* and *ROBO1*, respectively). Colour represents normalized gene expression (0-1). **(F)** Gene ontology (GO) enrichment plot illustrating GO terms upregulated in MS-associated ependymal cells. Note a variety of terms associated with cell adhesion (black boxes). Dot size represents the number of genes associated with a GO term, whereas dot colour represents adjusted p-value (p.adjust). **(G)** Heatmap visualizing the average expression of upregulated junction-associated genes and transporter-associated genes in MS donors (MS1-MS5) compared to control donors (C1-C4). Colour represents average gene expression.

### Ultrastructural evaluation of lesion-associated ependymal cells in MS provides further evidence that cell-cell adhesion is compromised

To determine whether multiciliated ependymal cell-cell adhesion was altered in MS as predicted based on our sequencing data, we completed ultrastructural evaluation of ependymal cells in MS that were adjacent to NAWM and lesions. In NAWM-adjacent regions, we observed normotypic cuboidal/low columnar ependymal epithelium with intact apically localized adherens junctions (Figure 4A). A classic dense network of astroglial processes was also found directly beneath the ependymal monolayer. In lesion-adjacent regions, ependymal cell-cell adhesion was disrupted (Figure 4B; Supp. Figure 2A). Although apical adherens junctions remained intact in most cases, the lateral membranes of lesion-adjacent ependymal cells, which are also bound by a variety of adhesion proteins such as cadherins,^26,39^ were separated by large, cavernous intercellular spaces, perhaps representative of CSF and/or interstitial fluid accumulation. In all cases of cell-cell “splitting”, we observed characteristic finger-like projections which protruded from adjacent lateral membranes towards one another, suggesting an attempt to re-establish cell-cell adhesion (Figure 4B; Supp. Figure 2A). The morphology of many lesion-adjacent ependymal cells was also altered, in that cells were irregularly shaped (non-cuboidal) or flattened (Figure 4C; Supp. Figure 2B). In general, MS ependymal cells appeared to accrue lysosomal debris (Figure 4A-B; Supp. Figure 2A-B), although this buildup did not seem to differ between NAWM-adjacent and lesion-adjacent regions. We did not observe any overt changes in the presence or structure of microvilli and cilia between NAWM-adjacent and lesion-adjacent ependymal cells (Figure 4A-B; Supp. Figure 2A-B). That said, cilia were typically not preserved well, so we cannot make a conclusive statement about ciliary alterations in MS based on these ultrastructural data. We noted the scattered presence of ciliary basal bodies in most ependymal cells (Figure 4A-B).

**Figure 4.**
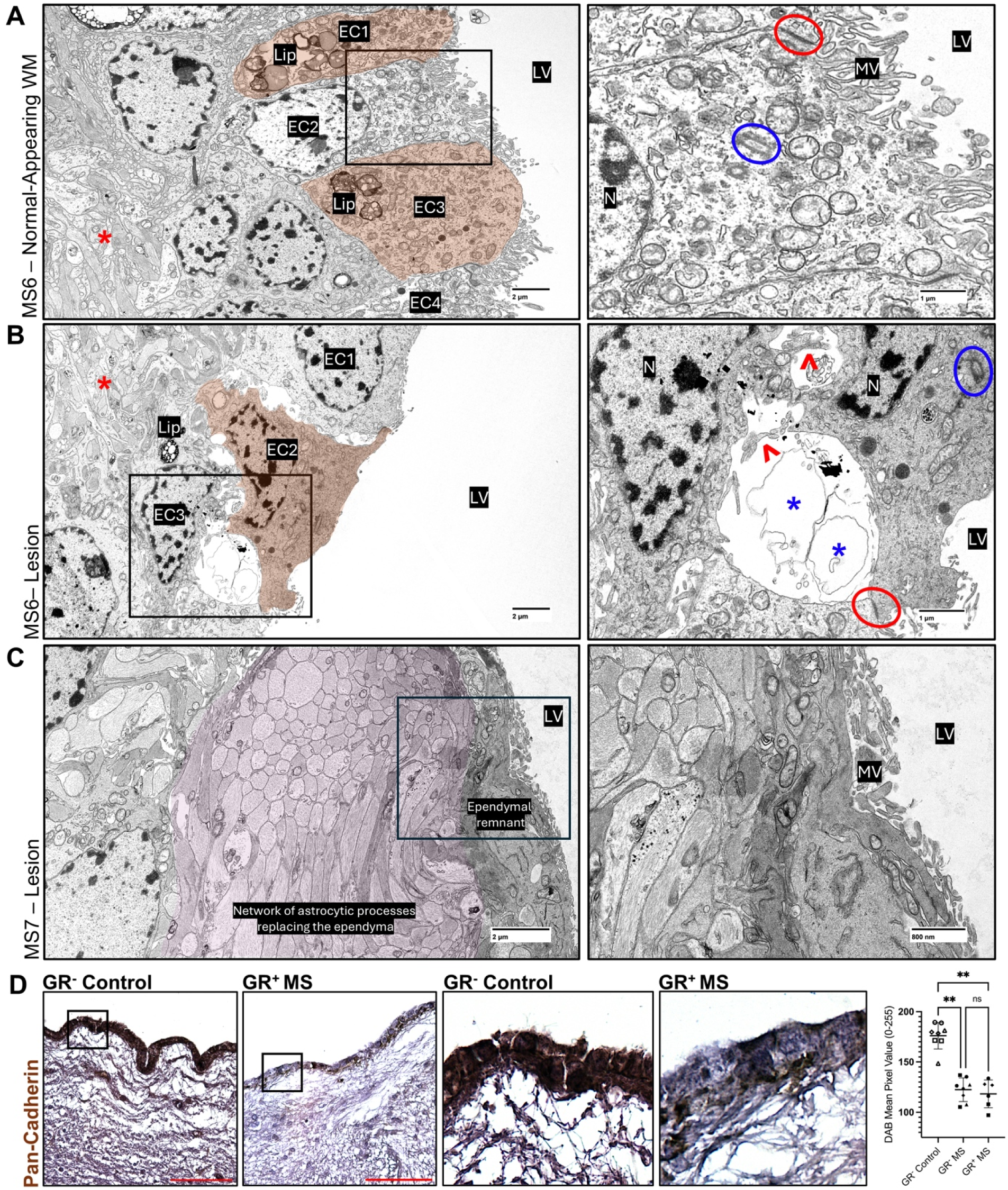
Ultrastructural evaluation of lesion-adjacent ependymal cells in MS provides further evidence that cell-cell adhesion is compromised. **(A)** (Left) Image of the ependymal lining in an MS brain (MS6), adjacent to normal-appearing white matter, demonstrating intact ependymal cell-cell adhesion. A black box indicates the region of the magnified panel on the right. A red star indicates subependymal astroglial processes. Scale bar = 2µM. (Right) Magnified image of lesion-distal ependymal cells with normotypic, intact cell-cell adhesion. A red circle denotes an adherens junction, and a blue circle denotes a ciliary basal body. Scale bar = 1µM. **(B)** (Left) Image of the ependymal lining in an MS brain (MS6), adjacent to a periventricular lesion, demonstrating disrupted ependymal cell-cell adhesion. A black box indicates the region of the magnified panel on the right. A red star indicates subependymal astroglial processes. Scale bar = 2µM. (Right) Magnified image of lesion-proximal ependymal cells with disrupted cell-cell adhesion. Red arrowheads point to lateral cellular processes ‘reaching out’ from detached ependymal cells and blue stars denote curious circular structures present in the space between separated ependymal cells, which we observed in all cases of cell adhesion disruption. A red circle denotes an adherens junctions and a blue circle denotes a ciliary basal body. Scale bar = 1µM. **(C)** (Left) Image an ependymal cell adjacent to a periventricular lesion (MS7) that has been completely flattened or resorbed (labeled ependymal remnant) and replaced by a dense ‘cobblestone’ network of invading astrocyte processes (purple-highlight). Scale bar = 2µM. (Right) Magnified image of ependymal remnant and astrocyte network. Scale bar = 800 nM. EC = ependymal cell; N = nucleus; MV = microvilli; LV = lateral ventricle; Lip = lipofuscin. **(D)** (Left) Representative unmagnified pan-cadherin images showing decreased staining intensity in ependymal cells lining GR+ blocks in MS compared to GR- blocks in control. Black boxes indicate the regions of the two magnified panels on the right. Scale bars = 200µM. (Right) Representative magnified images of ependymal pan-cadherin expression, and quantification of ependymal pan-cadherin (DAB) mean pixel value in GR- blocks in control compared to GR- and GR+ blocks in MS. This analysis showed a decrease in pan-cadherin (DAB) mean pixel value in ependymal cells lining GR+ and GR- blocks in MS compared to GR-blocks in control (p<0.01). Unfilled squares, triangles, circles, and diamonds represent individual control patients (H1, 2, 3 and 4, respectively; Supp. Tables 1-2). Filled triangles, squares, circles, and diamonds represent individual MS patients (MS1, 2, 3, and 4, respectively; Supp. Tables 1-2).

We also observed what could be interpreted as ‘intermediate’ stages of ependymal cell loss adjacent to periventricular lesions, where ependymal cells were somewhat intact, but subependymal astroglia had extended processes into the large intercellular spaces formed between separating ependymal cells (Supp. Figure 2B). In rare cases, we observed regions in lesion-adjacent tissue where ependymal cells had been completely flattened (Figure 4C). These regions were infiltrated by an abnormally dense network of astroglial processes, seemingly in an attempt to re-establish the functional properties afforded by an intact ependymal monolayer (Figure 4C), as has been described previously in hydrocephalic hyh mutant mice.^40^ The molecular mechanisms governing ependymal-astrocyte interactions in the maintenance of partial barrier capacity at the periventricular interface during neuroinflammation are clearly dynamic yet remain poorly understood.

Immunohistochemical analysis of cadherin proteins using a pan-cadherin antibody labeling most cadherin protein sub-families, demonstrated a decrease in DAB signal in the ependyma lining in both GR+ and GR- regions in MS compared to ependymal cells lining GR-regions in control (Figure 4D). In normotypic ependyma, we would expect homogenous expression of pan-cadherin throughout the monolayer (as seen in GR- regions in control), as cadherin proteins are expressed widely at ependymal cell-cell interfaces (both within and external to zonula adherens complexes).^26,39^ This observation could suggest that ependymal barrier alterations occur early in periventricular pathology pathogenesis in MS, prior to the development of *surface-in* gradients. This immunohistochemical observation was corroborated by our ultrastructural analysis, where cell-cell “splitting” was observed. Further, such features would also explain the clear upregulation of adhesion genes in our sequencing data. Indeed, upregulation of adhesion protein-associated genes of the armadillo/beta-catenin superfamily (*CTNNA3*, *CTNNA2*, and *CTNND2*), cadherin superfamily (*PCDH9*), and immunoglobulin superfamily (*CADM1*, *CADM2*) in MS ependymal cells (Figure 3G) might represent a compensatory mechanism to re-establish cell-cell contact. Similarly, changes in the expression of solute transporters (*SLC14A1*, *SLC1A2*) and genes governing cell shape and proliferation (*PRR16, SIPAL1, PTPN9*) (Supp. Figure 1A) may also represent a plastic ependymal cell response to disrupted epithelial polarity and/or cell-cell adhesion.

### Multiomic exploration of MS ependymal cells defines gene regulatory networks underpinning cellular reactivity and situates TNF and IFN***_γ_*** as candidate drivers of disease state

Our histologic, ultrastructural, and transcriptomic data suggested that MS ependymal cells enter a reactive state defined by cytoskeletal remodeling and altered cell-cell adhesion. To more precisely predict cell-intrinsic genetic regulation underpinning this MS-associated ependymal cell state, we utilized SCENIC+,^41^ a package that combines single-cell chromatin accessibility and gene expression data to identify candidate genomic enhancers and transcription factor (TF) binding motifs within said enhancers (Figure 5A). SCENIC+ then permits linkage of identified TFs to candidate enhancers and downstream targets genes and infers the direction of regulation (activating or repressing). Using this tool, we were able to construct GRNs that defined both control and MS-associated ependymal cell states (Figure 5A).

**Figure 5.**
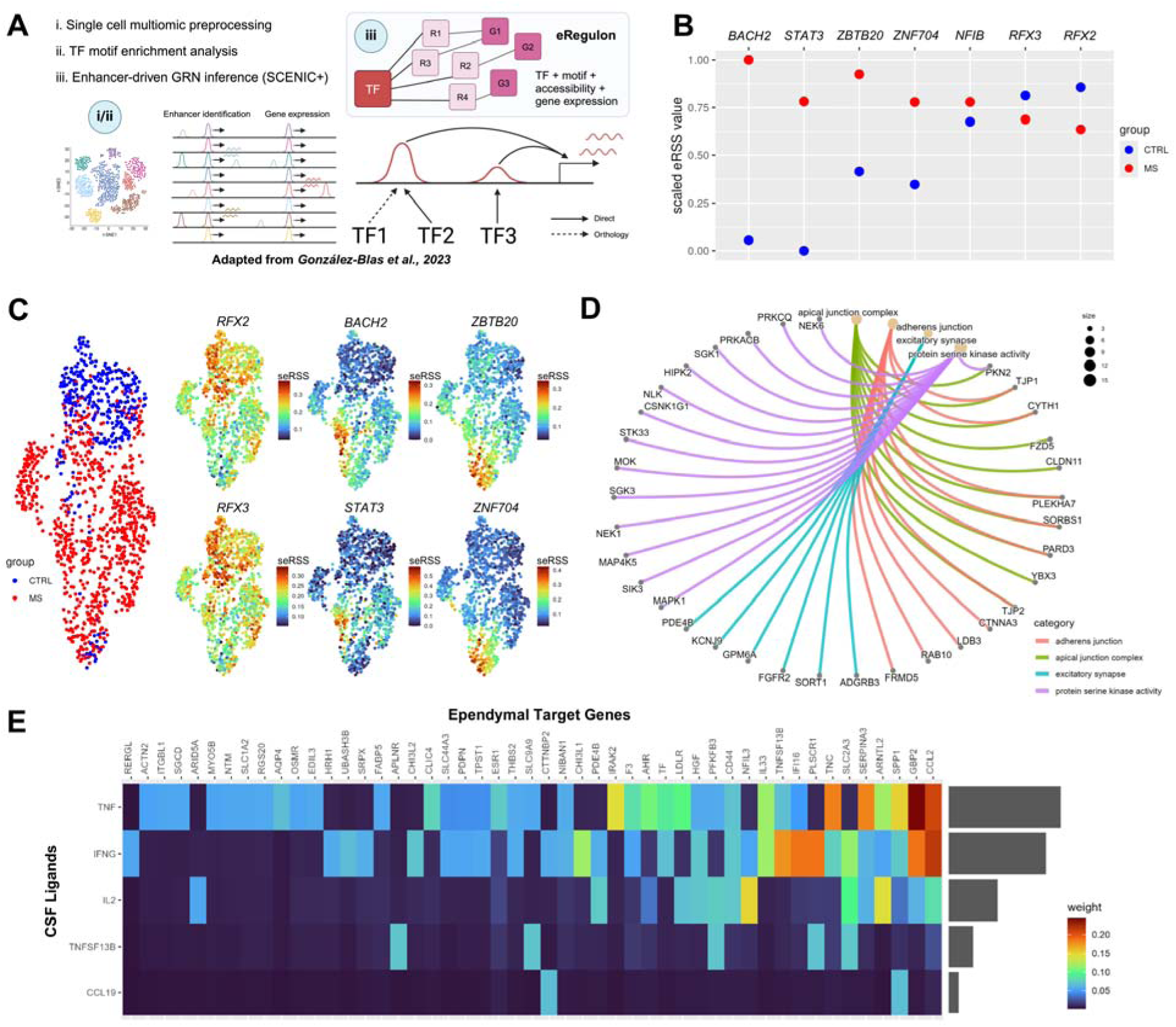
Multiomic exploration of MS ependymal cells defines gene regulatory networks underpinning cellular reactivity and situates TNF and IFN_γ_ as candidate drivers of disease state. **(A)** Schematic of the SCENIC+ workflow, adapted from González-Blas et al., 2023.^41^ **(B)** Dot plot representing scaled eRegulon Specific Scores (eRSS) for TFs preferentially acting in control (blue) and MS (red) ependymal cells. *BACH2*, *STAT3*, *ZBTB20*, and *ZNF704* showed significantly increased eRSS values in MS compared to control, whereas *RFX3* and *RFX2* showed significantly increased eRSS values in control compared to MS. **(C)** UMAP visualization of *RFX2*, *RFX3*, *BACH2*, *STAT3*, *ZBTB20*, and *ZNF704* scaled eRSS scores per cell, demonstrating unique enrichment of *BACH2*/*STAT3* in one population of MS (red) ependymal cells compared to control (blue) ependymal cells, and *ZBTB20*/*ZNF704* in another population of MS ependymal cells compared to control ependymal cells. Colour indicates scaled eRSS (seRSS) score (0-1). **(D)** Gene ontology plot showing the GO terms associated with the target genes of *BACH2*, *STAT3*, *ZBTB20*, and *ZNF704* that were enriched in MS ependymal cells compared to control ependymal cells. This analysis revealed that adherens junction-associated genes, synapse-associated genes, and serine kinase-associated genes are regulated by these TF networks in MS ependymal cells. Dot size indicates the number of genes associated with a GO term. Line colour indicates the GO term a specific gene is associated with. **(E)** Heatmap of ligands upregulated in MS CSF and correlated with periventricular damage, that were predicted to drive upregulated genes in MS ependymal cells. Colour indicates predicted interaction weight.

Plotting of scaled eRegulon specificity score (eRSS) values, which indicate the regulatory importance of a particular TF to a cell type or state of interest, allowed us to determine key TFs driving control and MS-associated ependymal cells states (Figure 5B). We filtered this list for TFs that acted as activators. This analysis uncovered *BACH2*, *STAT3*, *ZBTB20*, and *ZNF704* as the four primary TFs underpinning the MS-associated ependymal cell state, and *RFX3* and *RFX2* as key regulators of the control-associated ependymal cell state (Figure 5B). We then utilized our chromatin accessibility data (in isolation) to conduct TF footprinting and motif accessibility analysis using Signac^42^ and chromVAR^43^ packages, respectively, for the TFs with the top eRSS values in control and MS ependymal cells (*BACH2*, *STAT3*, *RFX2*, and *RFX3*; Supp. Figure 3A-B). We also profiled the basic gene expression of these TFs (Supp. Figure 3B). This allowed us to independently evaluate the results we generated using SCENIC+. We did not find a significant increase in motif accessibility for *STAT3*, *RFX3*, and *RFX2* (adj.p > 0.05) in control ependymal cells, whereas we did for *BACH2* (adj.p = 1.98E-12) (Supp. Figure 3B). Likewise, basic gene expression analysis of these same four TFs showed that only *RFX2* was significantly differentially expressed (adj.p = 6.93E-08) between control and MS ependymal cells (Supp. Figure 3C). These results stress the importance of integrating both chromatin accessibility data and gene expression data to make inferences about state-relevant TFs.

We next re-visited our SCENIC+ output and plotted the eRSS values of the four primary MS state-associated ependymal cell TFs on a UMAP of both control and MS ependymal cell populations to determine if the expression of these TFs was enriched in specific cell subpopulations. We found that *BACH2* and *STAT3* expression was unique to one population of MS-associated ependymal cells, which were in a more inflammation-associated state (IA-state), whereas *ZBTB20* and *ZNF704* expression was unique to a second (different) population of MS-associated ependymal cells, which were in a more regeneration-associated state (RA-state) (Figure 5C). This multiomic analysis therefore enabled us to uncover two disease-associated ependymal sub-states that both occur in MS and appear to be driven by entirely separate GRNs. When we conducted gene ontology analysis on the upregulated target genes of cells with high eRSS scores for *BACH2*/*STAT3* and *ZBTB20*/*ZNF704* in MS, we found that ependymal cells in both disease sub-states showed enrichment of genes associated with adherens junctions and synapse and neuronal guidance-associated factors (Figure 5D), as described previously in Figure 3. However, gene ontology analysis of MS ependymal cells governed more by *BACH2*/*STAT3* TFs showed unique enrichment of genes associated with immune regulation and inflammation (Supp. Figure 4A). Conversely, MS ependymal cells governed more by *ZBTB20*/*ZNF704* TFs showed unique enrichment of genes associated with gliogenesis, oligodendrocyte differentiation, and Wnt signaling (Supp. Figure 4A).

Finally, to predict what ligands might (upon binding) coax ependymal cells into the MS-associated state(s) we previously uncovered, we applied an interaction analysis pipeline using NicheNet.^44^ We filtered our list of potential regulatory ligands to those in the CSF of MS patients that positively correlate with the severity of periventricular pathology.^10^ That said, it is possible that ligands present in the inflamed periventricular parenchyma of MS patients (not the CSF) could also have an effect on ependymal cell function; for this reason, we also generated an unfiltered list of all predicted regulatory ligands and their ependymal target genes (Supp. Figure 4B). Our filtered analysis determined that the ligands with the highest predicted regulatory potential (that are also elevated in MS CSF) were TNFα, IFNγ, IL2, TNFSF13B, and CCL19. The latter 3 ligands regulated far fewer ependymal target genes than TNFα and IFNγ, and both TNFα and IFNγ were predicted to regulate shared ependymal target genes such as *CCL2* and *GBP2* (factors involved in immune regulation). Beyond these shared target genes, each ligand was also predicted to regulate distinct target genes, such as *IRAK2* (an interleukin-1 receptor-associated kinase 2, regulated by TNFα) and *SLC2A3* (a glucose transporter, regulated by IFNγ). Altogether, this interaction analysis situated TNFα and IFNγ as the two primary MS-enriched ligands which might coax normotypic ependymal cells towards an MS-associated state.

### Acute exposure of murine ependymal cells to IFN***_γ_*** in vivo increases barrier permeability, which is prevented by conditional knockout of ependymal interferon gamma receptor 1

To further test the impact of TNFα and IFNγ exposure on ependymal cells, we ran a small screen of these candidate ligands *in vitro* (Figure 6A). The screening assay suggested that IFNγ had a more deleterious effect on ependymal cells; 48h of exposure to IFNγ led to a significant reduction in the percentage of FOXJ1-positive cells compared to PBS control (p<0.001), whereas TNF did not (P>0.05) (Figure 6B). For this reason, we elected to focus on a single candidate ligand (IFNγ) for further exploration *in vivo*.

**Figure 6.**
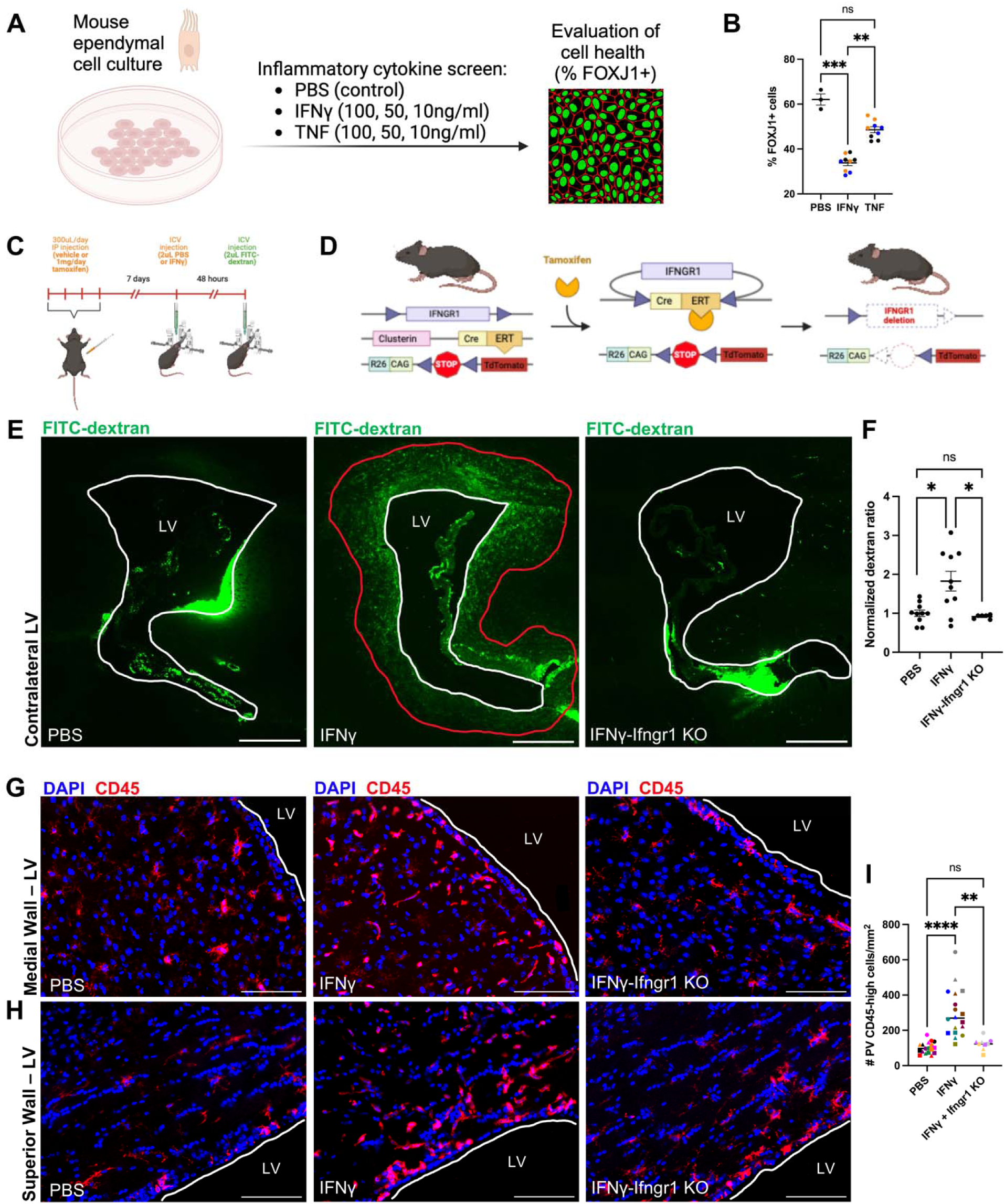
Acute exposure of murine ependymal cells to IFN_γ_ in vivo increases barrier permeability, which is prevented by conditional knockout of ependymal interferon gamma receptor 1. **(A)** Schematic of inflammatory cytokine screen used to determine the impact of IFNy and TNF on ependymal cell health. **(B)** Quantification demonstrating a significant decrease in the % of FOXJ1+ cells in response to IFNy compared to PBS control (p<0.001), but not in response to TNF (P>0.05), although the data was clearly trending toward a decrease. Black dots = 10nl/ml; Blue dots = 50nl/ml; orange dots = 100nl/ml. **(C)** Schematic of the surgical paradigm used to assess ependymal barrier permeability *in vivo*. **(D)** Schematic demonstrating the details of the Ifngr1^fl/fl^-Cre^ERT2^-lsl-Tdt mouse model. **(E)** Representative images of dextran infiltration from the lateral ventricle (contralateral to the site of injection) into the periventricular parenchyma in PBS-injected, IFNy-injected, and IFNy-injected + Ifngr1 knockout animals. LV = lateral ventricle. Scale bars = 100µM. **(F)** Quantification of normalized dextran ration indicating that IFNy increased dextran penetrance through the ependyma into the periventricular parenchyma compared to PBS sham (p<0.05) and that conditional knockout of ependymal Ifngr1 reversed this effect (p<0.05). **(G)** Representative images of periventricular CD45-positive cells adjacent to the medial wall of the lateral ventricle in PBS-injected, IFNγ-injected, and IFNγ-injected + Ifngr1 knockout animals. LV = lateral ventricle. Scale bars = 100µM. **(H)** Representative images of periventricular CD45-positive cells adjacent to the superior wall of the lateral ventricle in PBS-injected, IFNγ-injected, and IFNγ-injected + Ifngr1 knockout animals. LV = lateral ventricle. Scale bars = 100µM. **(I)** Quantification of the number of periventricular CD45-high cells/mm^2^, indicating that IFNγ injection increased the number of periventricular CD45-high cells/mm^2^ compared to PBS sham (p<0.0001), and that conditional knockout of ependymal Ifngr1 reversed this effect (p<0.01). Dot shape indicates ventricular region (filled circle = medial wall; filled square = lateral wall; filled triangle = superior wall), whereas dot colour represents individual animals. **p<0.01; ***p<0.001; ****p<0.0001

To test the hypothesis that IFNγ can drive the ependymal cell towards a reactive state in which cellular adhesion is compromised, we injected recombinant murine IFNγ intra-ventricularly to study the effects of acute exposure on ependymal barrier capacity (Figure 6C). In order to evaluate barrier capacity, we designed an assay in which a fluorescent dextran was injected intracerebroventricularly 24h following initial priming injection of IFNγ and permitted to circulate throughout the ventricular system for 20 mins (Figure 6C). Under normal homeostatic conditions, this 500 kDa dextran cannot pass through the ependyma into the periventricular parenchyma due to the expression of ependymal zonula adherens complexes and other junction-associated proteins.^26,39^ To evaluate the effect of IFNγ on ependymal cell-cell adhesion, we applied an inducible triple-transgenic line (Ifngr1^fl/fl^-Cre^ERT2^-lsl-Tdt) in which tamoxifen administration drives expression of cre-ERT2, and thus knockout of Ifngr1, only in cells expressing clusterin (Clu) (Supp. Figure 6A), which was ependymal cells and choroid plexus epithelial cells in the ventricular system (Figure 6D; Supp. Fig 6B). Clu-expressing cells also include a subset of a few other CNS cells, such as pial fibroblasts and astrocytes, but given our analysis was focused on the permeability of the ventricular wall in an acute experiment (with direct IFNγ injections into the ventricle) we do not anticipate this would impact our data. We found that acute exposure of mouse ependymal cells *in vivo* to IFNγ permitted increased periventricular dextran penetrance compared to PBS-injected control (p=0.015, as measured by a Kolmogorov-Smirnov test), suggesting that this cytokine can initiate a cellular response which drives dysregulation of ependymal cell-cell adhesion (Figure 6E). Conditional knockout of ependymal Ifngr1 led to a rescue of the barrier breakdown effects observed previously in response to IFNγ exposure (p = 0.016, as measured by a Kolmogorov-Smirnov test), confirming that the altered permeability of the ventricular wall was regulated by cell autonomous mechanisms (Figure 6E). We were also interested in the effects of increased ependymal interface permeability on the periventricular parenchyma following IFNγ exposure. We assessed the presence of periventricular CD45-high cells (immune cells and activated microglia) following exposure to IFNγ both with and without ependymal-specific knockout of Ifngr1. We found that IFNγ stimulated a large increase in periventricular CD45-high cells compared to PBS-injected control, and that ependymal-specific knockout of Ifngr1 rescued this effect (Figure 6G and H). In summary, these data confirmed that binding of IFNγ to ependymal cells led to a reactive response which increased the permeability of the ependymal interface and then presumably led to an inflammatory periventricular response.

## Discussion

Although there is consensus that the periventricular region of the MS brain is consistently subject to pathology,^8,45,46^ and that damage in this region correlates well with severity of disease progression,^4,11,46–51^ there are few hypotheses as to why this is the case. What is known is that periventricular pathology exists in a distinct *surface-in* gradient pattern,^6,8^ suggesting that CSF contributes to the pathogenesis of this damage. Our work supports this notion by revealing a striking correlation between the location of ependymal cell pathology and regions of periventricular *surface-in* gradients in MS. We define the gene regulatory networks responsible for mediating a shift of normotypic ependymal cells towards an MS-associated state and provide histologic, ultrastructural, and transcriptomic/epigenomic evidence that widespread alterations to ependymal cell adhesion occur in MS. We suggest that these changes to the ependymal monolayer result in increased permeability of the ventricular wall, which may play a role in the emergence of periventricular pathology. We computationally predict which ligands, upon binding to ependymal cells, may drive ependymal cells into a reactive state, and prove that IFNγ alone can modulate ependymal permeability *in vivo*. Through our analysis of the involvement of ependymal cell dysregulation in MS disease pathogenesis, we highlight an important relationship between disease-specific repertoires of CSF ligands, their influence on ependymal cell function, and the resultant impact on the pathological/inflammatory status of the periventricular region.

There are few infiltrating immune cells in periventricular regions in MS,^52^ suggesting that diffusion of soluble factors from the CSF may be central to disease pathogenesis at this brain border. If toxic CSF solutes are drivers of periventricular *surface-in* gradients, we speculate that ependymal cells are the first to be impacted, before the onset of widespread periventricular damage. This is consistent with our observation that regions of bona fide ependymal cell pathology (i.e., GFAP and vimentin upregulation) could predict regions of periventricular *surface-in* gradients of demyelination and neurodegeneration. Conversely, regions with less developed ependymal cell pathology (i.e., vimentin upregulation only) did not predict this damage. Similarly, ependymal pan-cadherin alteration was also seen in both GR- and GR+ regions in MS, further suggesting that there is early disruption of the ependyma in periventricular MS pathogenesis. Related observations have been made in a murine model of CNS demyelination, wherein early infection of ependymal cells by Theiler’s encephalomyelitis virus is followed by periventricular demyelination, axonal damage, and infection of microglia and oligodendrocytes.^53^ TLT-containing models of MS such as in the adoptive transfer (A/T) model of PLP-Th17 cells in SJL/J mice,^54–56^ show that meningeal inflammation is associated with a disrupted glial limitans superficialis, subpial demyelination, microgliosis, and axonal loss, particularly in older mice.^56^ This further suggests that inflammatory factors in the CSF mediate peri-brain border damage.

Although it is now clear that both murine and human ependymal cells are highly reactive under neuroinflammatory conditions, it is unclear why ependymal pathology and associated *surface-in* gradient pathology is found in defined periventricular locations in MS, scattered throughout the ventricles. Studies of the aging human brain show that the ventricular horns undergo increased loading states (i.e., stretching of ependymal cells) and are most susceptible to developing periventricular white matter pathology.^32,57^ Periventricular regions abutting the ventricular horns have also shown unique susceptibility to pathology in MS,^7^ suggesting that the loading state of the ventricular wall may influence where ependymal and periventricular pathology exists in MS. An additional variable that could influence this proposed disease model is the location from which proinflammatory solutes enter the ventricular system (i.e., blood-brain interstitium-CSF, blood-meninges-CSF, blood-choroid plexus-CSF). For example, cases of MS in which there is pronounced choroid plexus inflammation associate with more extensive periventricular pathology.^24^ How MS-associated cytotoxic factors exit the choroid and enter the CSF, where they can theoretically initiate and/or exacerbate periventricular pathology, is still an area of nascent inquiry.

Ependymal cell intrinsic factors could also explain heterogeneity in periventricular pathology emergence. A previous study demonstrated that mammalian ependymal cilia exist in discrete modules, where they orchestrate regionally localized patterns of CSF microcirculation that can preferentially distribute molecular cues throughout the CNS.^58^ Cytotoxic factors that have entered the CSF may therefore be confined to specific regions of the ventricles as a result of local cilia-mediated flow networks. A study in EAE showed that disrupted planar cell polarity, which is critical for proper motile ciliary beating, develops under autoimmune inflammatory conditions^36^. Whether this is a contributing factor to the heterogeneity of periventricular damage seen in human disease in not clear. Alternatively, perhaps variability in the ependymal junctional protein repertoire across the ventricles plays a role in regulating region-dependent periventricular pathology.^29,39^ Ependymal cells typically maintain a collection of adherens (e.g., ß-catenin, cadherins) and tight junction-associated proteins (e.g., zonula occludens family: Zo1/*Tjp1* and Zo2/*Tjp2*), although complete tight junctions are notoriously rare and located within specific populations lining circumventricular organs.^39^ Though ependymal ciliary heterogeneity is widely acknowledged,^29,59^ a thorough census of ependymal junctional protein expression across the ventricular system is lacking in mice and humans^29^ and could contribute to a greater understanding of their regulatory influence on ventricular wall permeability.

Beyond the influence of innate heterogeneity in ependymal cell function, there could also be an element of selective vulnerability in response to cytotoxic CSF factors, where certain ependymal cells are more likely to become damaged. A recent study found that a specific population of ependymal cells in the midbrain is highly immune responsive in aging, upregulating a variety of genes associated with a response to interferons and major histocompatibility complex class I regulation.^60^ Thus, inflammatory regulation in specific ependymal cell populations could further diversify CSF flow patterns and/or junctional protein stability, resulting in defined periventricular regions becoming more damaged. This hypothesis is further supported by our finding that ependymal cells exist in two disease-associated substates in MS, which are mediated by separate gene regulatory networks. Cells in one state appeared to be more immune- and inflammation-responsive (*BACH2*/*STAT3*-associated) than cells in the second state (*ZBT20*/*ZNF704*-associated), although both populations showed similar dysregulation of junctional protein-associated genes. Interestingly, the *ZBT20*/*ZNF704*-associated population uniquely upregulated a variety of genes associated with the Wnt-signaling pathway, which when activated, is known to induce ß-catenin signaling and substantiate cellular barrier properties.^61^ Perhaps this population is turning on a compensatory regenerative program to help re-establish normal permeability at the ventricular interface. This proposition is further supported by the fact that the *ZBT20*/*ZNF704*-associated population upregulated genes associated with CNS development, glial cell differentiation, and oligodendrocyte maturation. While brain ependymal cells are thought to lack stem cell potential,^26,34,62^ there is latent accessibility of an oligodendrocyte-associated genetic program in spinal cord ependymal cells, which is thought to permit generation of ependymal-derived oligodendrocytes following spinal cord injury.^63^ Perhaps this genetic program is serving an alternative, cell-intrinsic purpose to promote cellular regeneration in some brain ependymal cells in response to neuroinflammation.

In addition to characterizing the reactive ependymal cell state at the transcriptomic and epigenomic level, we also predicted ligands which might drive cellular reactivity and alterations to ependymal cell barrier properties that we observed in human tissue. We found that TNF and/or IFNγ were predicted to regulate a variety of target genes in ependymal cells associated with immune responsivity (e.g., *CCL2*, *GBP2*, and *SERPINA3*) and cell attachment and transport (*SPP1*, *SLC2A3*, and *CD44*). Of note, *SERPINA3*, the human orthologue of *Serpina3n*, is a well-established disease-associated astrocyte and oligodendrocyte marker gene^64–67^ that is also elevated in the CSF of patients with progressive MS.^68^ We recently demonstrated that the reactive ependymal cell transcriptome in MOG-EAE is strikingly similar to reactive astrocytes, including upregulation of *Serpina3n*, and a variety of other interferon response genes.^33^ Others have revealed similar dysregulation in EAE, such as ependymal expression of *Irgm1* (an interferon inducible GTPase) and genes encoding chemokines such as *Ccl20*, akin to our finding that *CCL2* is upregulated in MS ependymal cells under the control of both TNF and IFNy.^69^ Together, these data demonstrate consistency in mouse and human reactive ependymal cell responses to central nervous system autoimmunity, and hint at the more general importance of serine protease inhibitors in mediating reactive glial responses following CNS insult.

While both TNF and IFNγ were predicted to modulate ependymal target genes in MS, an *in vitro* screen suggested that IFNγ might have a more pronounced acute effect on cell health than TNF, so we focused on assessing the impact of this ligand *in vivo*. We found that IFNγ could initiate changes to ependymal cell barrier properties, as represented by increased penetrance of a fluorescent dextran from the ventricles, through the ependyma, and into the periventricular parenchyma. Although a novel finding at the ependymal interface, IFNγ-mediated disruption of gut epithelial cell barrier properties is well-described.^70–72^ IFNγ can induce a decrease in junctional protein expression in intestinal epithelial cells, which is associated with elevated JAK/STAT pathway activation and serine protein kinase activity;^71^ given we found that these same pathways were activated in ependymal cells in MS, it is likely that similar mechanisms are regulating ependymal cell adhesion alterations during CNS neuroinflammation. We also found that increased periventricular dextran penetrance was associated with increased presence of periventricular CD45-positive cells; we interpreted this to signify that IFNγ had increased access to the periventricular region due to increased ependymal permeability, where it could activate resident microglia. While this is merely a hypothesis, we showed that ependymal-specific knockout of Ifngr1 prevented barrier leakiness and suggest that this reduced the amount of IFNγ entering the periventricular region, and thus the increased presence of CD45-positive cells (which we suspect are primarily activated microglia, given the acute nature of the experiment). Alternatively, and/or concomitantly, ependymal cells could secrete proinflammatory cytokines in response to IFNγ binding, which could theoretically promote periventricular glial reactivity.

In summary, we employed ultra-high-field MRI-guided immunohistochemistry, electron microscopy, and multiomic single nucleus RNA/ATAC sequencing to deeply phenotype human ependymal cells in MS. Our data revealed that ependymal cell pathology is a direct correlate of periventricular *surface-in* gradients of pathology in MS. Ependymal reactivity can be initiated by inflammatory cytokines in the CSF and is associated with striking remodeling of ventricular wall permeability. These data suggest that there is an important relationship between the content of CSF, ependymal cell health, and the pathological/inflammatory status of the periventricular region. Future work could explore therapeutically modulating the periventricular milieu. This could help uncover how proinflammatory ligand binding initiates ependymal barrier property remodeling, and how this ependymal specific response might trigger periventricular glial reactivity. Further, elucidating mechanisms of CSF-mediated ependymal cell dysregulation may contribute to an improved understanding of VZ-SVZ alterations in aging and disease.

## Method

### Magnetic resonance imaging

In collaboration with the Douglas Bell Canada Brain Bank (DBCBB), 4 human MS brains (n=3 male; n=1 female) and 4 age- and sex-matched non neurological disease control brains were selected for study. Each brain was separated longitudinally with one hemisphere cut into coronal sections from frontal, middle, and caudal regions of the brain and preserved in 10% neutral buffered formalin. Each of these 23 total brain slabs (n = 12 MS; n = 11 control) were scanned overnight for 17 hours with a Siemens 7 Tesla Human MRI system (MAGNETOM Terra) using a one channel transmit and 32 channel receive (1Tx/32Rx) head coil (Nova Medical). Acquisitions included a 3D Multi-Echo GRE (ME-GRE) sequence of 0.32mm isotropic resolution (TR = 46ms, TE = [6.84, 11.57, 16.30, 21.03, 25.76, 30.49]ms, readout flip angle = 15°, 25 averages) and a 3D MP2RAGE sequence of 0.32×0.32×0.64mm spatial resolution (TR/TE = 3000/2.08ms, Turbo Factor = 24, echo spacing = 6.6ms, readout flip angle 1/2 = 8/8°, TI1/TI2 = 183/900ms, 25 averages). The 3D MP2RAGE sequence was used to compute a map of tissue longitudinal relaxation time (T1), sensitive to myelin, neuronal, and axonal densities, and the ME-GRE sequence was used to compute a map of tissue apparent transverse relaxation time (T2*), sensitive to myelin density and iron deposition.

Specifically, for each 3D MP2RAGE measurement, a map of T1 was computed using the Siemens MP2RAGE-based T1 fitting algorithm onsite. Individual T1 maps and ME-GRE magnitude images were averaged to yield T1 maps and ME-GRE images with high signal-to-noise ratio (SNR) for each brain slab. The averaged T1 map was then upsampled in resolution and registered to the averaged ME-GRE magnitude image using FLIRT (FSL, v7.2). Based on averaged ME-GRE magnitude data, a map of T2* was computed using a voxel-wise non-linear, least-squares fitting approach in MATLAB (R2020b, MathWorks). Finally, to extract tissue-specific T1 and T2* maps, the first echo image of the averaged ME-GRE magnitude image was used to segment grey matter and white matter, via the ilastik Pixel Classification workflow.^54^

Once scanned, each slab was subdivided into angular segments surrounding the ventricle and geodesic bands from the ventricle to the meninges (**Figure 1A**). The segmentation of each slab into angular bands, each of 15° coverage, was performed in a counterclockwise fashion, starting on the superior-inferior brain axis, and going from the superior to inferior and medial to lateral directions, for a total of 24 bands (numbered 1 to 24 in order). Segmentation of tissue into geodesic bands was performed successively from the ventricular surface towards the cortical surface, with a consistent band width of 1.5mm. The range of geodesic bands for calculating total change in T1 and T2* was chosen so that the first band was second to the ventricular surface (starting at 1.5mm distance from ependyma), to avoid partial volume effects from CSF voxels, and the last band was second to the boundary of subcortical white matter with cortical gray matter to avoid partial volume effects from grey matter. T1 and T2* relaxation time values were subsequently extracted for each segment and averaged voxel-wise through the depth of each slab. Plots of the change in relaxation times along concentric geodesic bands were generated for each angular segment. Angular segments that showed relatively consistent T1 and T2* relaxation times along geodesic bands were identified as gradient negative (GR^-^) while angular segments that showed a peak in relaxation times near the ventricle that decreased and levelled off with increasing distance, in addition to meeting specific criteria (see Results section), were identified as gradient positive (GR^+^) regions. With these regions identified, sections of tissue both positive and negative for gradients were selected and resected. 28 blocks from MS brains (n = 16 GR+; n = 12 GR-) and 18 control (n = 2 GR+; n = 16 GR-) were processed and dehydrated, embedded in paraffin, sectioned (5um) and mounted on Superfrost^TM^ Plus slides in preparation for histology.

### Hematoxylin & Eosin (H&E)

Slides were deparaffinized by washing in xylene for 10 minutes followed by another xylene wash, two washes each with 100% and 95% ethanol followed by 70% ethanol, all for 5 minutes. After washing in water, the slides were then soaked in hematoxylin for 3 minutes before being dipped in 1% HCL in 70% alcohol and 2% ammonium hydroxide in 70% alcohol, then 3 minutes in eosin with washes under running water between each step. The tissue was then dehydrated for 3 minutes each in 70%, 95%, 100% ethanol and xylene before mounting with permount toluene solution. Once complete, each H&E-stained slide was evaluated to determine the location of the ependyma. To be used for downstream histological stains and quantification, the ventricular wall had to be bounded by a ciliated, simple layer of cuboidal/low columnar ependymal epithelium. Regions that were entirely devoid of ependymal cells were not analyzed as postmortem stripping of the ventricular wall (associated with autopsy) could not be ruled out as a cause.

### Luxol Fast Blue (LFB)

Slides were deparaffinized and rehydrated in xylene for 5 minutes, followed by 10 dips in 2 more xylene baths, and 15 dips each in 2x 100% and 2x 95% baths. For staining, slides were immersed in a 0.2% Luxol Fast Blue solution at 60C for 3 hours. Excess stain was removed with 95% ethanol and washing in distilled water. Slides were then washed in 0.05% lithium carbonate and twice in 70% ethanol for 10-20 seconds to differentiate the stain. Next, they were washed in distilled water for 5 minutes and stained with a working cresyl violet solution for 1 minute. Slides then underwent 2x washes in 95% ethanol, one in terpineol-xylene solution, and one in xylene for 5 minutes each before mounting with Prisma B coverslipper.

### Bielschowsky stain

Slides were deparaffinized as described for H&E then placed in distilled water. Slides were then transferred to a pre-warmed 37C 20% silver nitrate (Fischer scientific, cat # S181-500) solution for 15 minutes. The slides were removed and placed in distilled water while 28% ammonium hydroxide (ACP, cat # A-6320) was added drop by drop until the solution turned clear. Once clear, two more drops were added to make ammoniacal silver. The solution was then transferred into a new pre-warmed coplin jar, and the slides were added and incubated at 37C for an additional 15 minutes. After, slides were placed in a 1% ammonium hydroxide solution for 1-3 minutes and 10-25 drops of developing solution (10% formalin ACP, cat # F-5940; nitric acid Anachemia, cat # AC-6525; citric acid Anachemia, cat # AC-2514) was added to the ammoniacal silver, mixing carefully. Slides were returned to the developing ammoniacal silver solution for 3-5 minutes periodically checking for black neurofibrils under the microscope. Once visible, the reaction was stopped by placing slides in 1% ammonium hydroxide for 3 minutes followed by a 5-minute incubation in 5% sodium thiosulfate (A&C, cat # S-440). The slides were then rinsed with distilled water 3 times and dehydrated in graded alcohol (70%, 95% and 100%) for 5 minutes each. Slides were mounted with permount mounting medium.

### Immunohistochemistry (IHC)

Human FFPE tissue blocks were sectioned at 7 um thickness using a microtome. A 40-45C hot water bath was used to stretch the tissue sections, and glass slides were used to mount them. The sections were dried overnight in an oven at 37C. Slides were baked at 40C overnight and deparaffinized as described above for H&E with an additional 20-minute incubation in 0.3% hydrogen peroxide in methanol for chromogenic IHC. For antigen retrieval, slides were washed in phosphate-buffered saline (PBS) and incubated in 10 mM sodium citrate buffer pH 6 at 95C for 20 mins. Sections were washed in PBS and blocked using 0.5% triton-X (Sigma-Aldrich, cat # T9284) in 10% goat serum for 30 minutes at room temperature. Primary antibodies were diluted in Normal Antibody Diluent (BD09, Immunologic) and incubated overnight at RT. These included: mouse anti-GFAP (1:400; ab190288, Abcam), rat anti-vimentin (1:100; MAB2015, Novus Biologicals), rabbit anti-Iba1 (1:1000; 17198S, Cell Signaling Technology), and rabbit pan-cadherin (1:100; ab51034, abcam). Following primary antibody incubation, slides were washed with PBS and monoclonal antibody-exposed slides incubated in Post Antibody Blocking BrightVision Solution 1 (DPVB-HRP, Immunologic) diluted 1:1 in PBS for 15 minutes. After another wash, all sections were then incubated in BrightVision Poly-HRP-Anti Ms/Rb/Rt IgG Biotin-free Solution 2 (DPVB-AP, Immunologic) diluted 1:1 in PBS for 30 minutes. Following another wash in PBS, slides were incubated in a 3,3’-diaminobenzidine (DAB) solution (SK-4100, Vector Laboratories) for 5 to 7 minutes. This reaction was stopped by putting slides in PBS followed by washing in demi-water. Slides were counterstained with hematoxylin for 30 seconds and washed under running water for 5 minutes. Finally, slides were dehydrated in graded alcohol (70%, 95%, 100%) and xylene for 5 minutes each before mounting with vectamount (H-5000, Vector Laboratories).

Following cardiac perfusion with 1x PBS at RT and pre-chilled 4% PFA, mouse brains were harvested and post-fixed in 4% PFA for 2–12 h at 4°C (n=10 PBS; n=10 IFNγ; n= IFNγ-Ifngr1 KO). Brains were immersed in 30% sucrose at 4°C until they sank (24–72 h) and snap-frozen in O.C.T. (14-373-65, Andwin Scientific Tissue-Tek©) and stored at −80°C. Tissue was coronally cryo-sectioned at 15 μm on Superfrost Plus slides (22-037-246, Fisher Scientific) and stored at −80°C. Sections were blocked with 0.5% Triton-X and 10% donkey serum for 2h at RT. Sections were incubated with primary antibodies in 0.5% Triton-X (in 10% donkey serum) overnight at 4°C: rat anti-CD45 (1:50; 550539, BD Pharmingen™) and rabbit anti-Iba1 (1:1000; ab178846, Abcam). Sections were washed twice with 1x PBS and incubated with Alexa Fluor secondary antibodies in the dark for 2h at RT (1:200). Following two washes with 1x PBS, sections were incubated in the dark with DAPI (1:5000; D1306; Invitrogen) for 10 min at RT. Sections were washed twice more with 1x PBS before being mounted (PermaFluor™ Mounting, TA030FM, Fisher Scientific) and cover-slipped (Micro cover glass, VWR).

### Image Analysis and Statistics

Human slides were imaged using a light microscope at 40x and a Leica TCS SP8 Multiphoton Confocal microscope at 40x magnification, in oil. Histological brightfield images were loaded into FIJI (ImageJ) for cell counts (Iba1), axon analysis (Bielschowsky), and DAB intensity quantification (pan-cadherin). Specifically, DAB intensity was calculated by dividing the colors of chromogenic images into green, purple, and brown images using the color deconvolution tool with the ‘H DAB’ vector selected. Brown images were thresholded and, using the freehand selection tool, the ependymal monolayer was selected for analysis of DAB mean pixel value intensity (0-220). Fluorescent images were similarly loaded into FIJI (ImageJ) for cell counts (GFAP) and corrected total cell fluorescence (CTCF) analysis (vimentin). To conduct CTCF analysis, manual cell segmentations of at least 20 ependymal cells were repeated three separate times to obtain an average total cell area, mean intensity, and integrated density. Background selections of the same area were taken for comparison. Values were input into a CTCF equation to generate individual CTCF metrics: CTCF□=□integrated density□−□(area of selected cell□×□mean fluorescence of background readings). Mouse slides and culture plates were imaged using a Zeiss Axio Observer epifluorescence microscope at 20x magnification. Fluorescent images were loaded into FIJI (ImageJ) for cell counts (FOXJ1, CD45) and dextran ratio quantification. Appropriate statistical tests were run and graphs generated using GraphPad Prism v9.0.

### Single nucleus RNA sequencing of archived frozen human periventricular brain tissue

The periventricular region adjacent to the lateral ventricle was dissected en-bloc from two coronal fresh-frozen human brain slabs per patient sourced from the Douglas Bell Canada Brain Bank (n=4 control patients [3F, 1M], n=8 total brain slabs; n=5 MS patients [2F, 3M], n=10 total MS brain slabs) (Supp. Table 3). Of the two brain slabs sourced per patient, one was more rostral, and the other caudal, although perfect rostral-caudal standardization could not be completed between patients given limited access to archived human brain tissue. This limited access to tissue also left us underpowered to assess sex differences. Nevertheless, this design still permitted us to obtain representation of the ventricular lining from two separate periventricular brain regions per patient. Directly prior to nucleus isolation, frozen brain tissue was removed from the −80 and transferred to dry ice, where the ventricular lining was further micro-dissected to enrich for ependymal cells. Importantly, we dissected periventricular white matter and overlying ependymal cells from each sample such that the sequencing data would be comparable to our MRI-guided histological analysis, which focused exclusively on gradients in periventricular white matter, not gray matter.

Processing of periventricular samples was pooled, in that two control (1F, 1M) and two MS (1F, 1M) samples were processed together, respectively, until all samples had been completed. We used this design so that samples could be demultiplexed later during the sequencing pipeline, and to limit the influence of batch effects on specific experimental groups or sexes. Each individual sample was weighed immediately following dissection on dry ice; sample weights ranged from 25-45mg, depending on the accuracy of dissection. We prioritized immediate processing following dissection to retain RNA quality, rather than complete standardization of sample weight. This meant that each pooled sample had an approximate weight of 50-90mg. Processing samples >100mg with our protocol led to increased debris and clogging during droplet generation, so we elected to stay beneath that threshold.

Nuclei extraction was conducted using a modified protocol adapted from Maitra & colleagues.^73^ All steps were completed either on ice or at 4 degrees in a pre-cooled centrifuge. All buffers described in the protocol were prepared as described previously^73^ and were pre-chilled at 4 degrees before use. Pooled samples were carefully dropped into a douncer (357542, Wheaton) that was pre-filled with 750µL of lysis buffer. Another 750µL of lysis buffer was then added to coax any tissue that had stuck to the sides of the douncer into the lysis buffer below. Tissue was then carefully dounced with a loose pestle 10 times, and a tight pestle 5 times. When pulling the pestle up from the bottom of the douncer, it was important to keep it below the surface of the solution to avoid addition of air bubbles that damaged nuclei. If the mixture looked heterogeneous after initial douncing, this process was continued with a tight pestle until the solution appeared homogenous. Dounced solutions (1.5mL) were then transferred to a 15mL falcon tube, mixed with 3.5mL of lysis buffer, and left to incubate on ice for 5 mins to complete cell lysis. 5mL of wash buffer was then added to stop the lysis reaction and nuclei solutions were passed through a 30µM strainer (130-098-458, Miltenyi Biotech) to remove debris. Nuclei were then centrifuged at 500g for 5 mins at 4 degrees. The supernatant was subsequently decanted without disrupting the pellet, and 5mL of fresh wash buffer was added, mixing 8-10 times with a serological pipette to resuspend the nuclei. Samples were then taken through 2 more successive rounds of filtration and centrifugation as described. After the third filtration/centrifugation, nuclei were resuspended with 5mL of wash buffer, not passed through a strainer, and centrifuged at 500g for 5mins at 4 degrees. The supernatant was decanted, and this time nuclei were resuspended with 3.1mL of wash buffer plus 900uL of chilled debris removal solution (130-109-398, Miltenyi Biotech). This mix was carefully overlaid with 4mL of wash buffer and centrifuged at 3000g for 10 mins at 4 degrees. The top two phases were then aspirated and discarded, and the bottom phase (containing the nuclei) was mixed with additional wash buffer to a final volume of 15mL. The falcon tube was inverted 2-3x and centrifuged at 1000g for 10 mins at 4 degrees. The final supernatant was aspirated completely, and nuclei were resuspended using low-bind pipette tips in an appropriate volume of buffer for single nucleus RNA sequencing. A few nuclei aliquots from each sample were set aside for staining with DRAQ5 (65-0880-92, ThermoFisher) to assess nuclei for blebbing or other indications of low quality using a 40x objective, and for nuclei counting using a hemocytometer and an Attune NxT Flow Cytometer, Model AFC2 (A29009, Invitrogen).

Single nucleus RNA sequencing was completed as per the Chromium Next GEM Single Cell Multiome ATAC + Gene Expression protocol (CG000338, Rev E to Rev F). In brief, this included transposition, GEM generation and barcoding, post-GEM cleanup, pre-amplification PCR, ATAC library construction, cDNA amplification, and gene expression library construction. The quality of both ATAC and RNA libraries was evaluated at Genome Quebec using a LabChip® GX Touch™ nucleic acid analyzer system. Multiome libraries were sequenced using an Illumina NovaSeq 6000 instrument at a sequencing depth and read length recommended by the Multiome protocol (CG000338).

### Sequencing data pre-processing

Demultiplexing, genome alignment, gene quantification, and peak accessibility in single nuclei were performed with Cell Ranger ARC 50. Specifically, data were processed using the cellranger-arc count v.2.0.2 and aligned against the human genome (GRCh38) to generate count matrices for the RNA and ATAC modalities pipeline (v.7.0.1).^74^ Given two patients were merged per bead, and thus fastq files were multiplexed, BAM files generated from the cellranger-arc count were demultiplexed through SNP information of the sample using 1000 Genome based common variants (genome1K.phase3.SNP_AF5e2.chr1toX.hg38.vcf). The subsequent data processing, normalization, and batch integration on both assays were done using Seurat^75^ (v.5.0.1) and Signac^42^ (v.1.12.0) in R (v.4.1.2).

### Single nucleus RNAseq analysis

Gene expression data were processed using Seurat (v5.0.0). Raw counts were normalized with the LogNormalize method (scaling factor = 10,000), and the top 2,000 highly variable genes were identified using a variance-stabilizing transformation (VST). Data were scaled, and principal component analysis (PCA) was performed using these variable features. The first 20 principal components were used to construct a k-nearest neighbor (KNN) graph, followed by clustering with the Louvain algorithm (resolution = 0.4). Uniform Manifold Approximation and Projection (UMAP) was applied for dimensionality reduction and cluster visualization, with clusters displayed based on their seurat_clusters identity. Differentially expressed genes (DEGs) between clusters was calculated using the Wilcox test implemented in Seurat. DEGs were selected using an adjusted p-value of less than 0.05 and a log2 fold-change threshold of 0.25. Gene Ontology (GO) analysis of DEGs was performed using clusterProfiler.^76^

### Single nucleus ATACseq analysis

Chromatin accessibility data were analyzed using Signac (v1.12.0). The ATAC assay was set as the default, and peak features with a minimum count of 50 were retained. Term Frequency – Inverse Document Frequency (TF-IDF) normalization was applied to the filtered features. Latent Semantic Indexing (LSI) was performed on the top 50 components, followed by dimensionality reduction using Uniform Manifold Approximation and Projection (UMAP) with dimensions 2 to 30. A k-nearest neighbor (KNN) graph was constructed using the LSI reduction, and clustering was performed using the Louvain algorithm with the Leiden refinement (algorithm = 3). Cluster determination was guided by an elbow plot of LSI components and depth correlation analysis.

### Multimodal data integration

The filtered_feature_bc_matrix.h5 file which is one of the output files of the cellranger-arc pipeline and contains both metrices for transcriptome and chromatin accessibility was used as the input file. The following criteria were used to process the samples and remove low quality nuclei: nFeature_RNA > 1000, nFeature_RNA < 7500, percent.mt < 10, nFeature_ATAC > 200, nFeature_ATAC < 5000, TSS.enrichment > 2, nucleosome_signal < 4. The contaminated RNA content of the nuclei was removed using the SoupX package.^77^ To correct batch effects between individual samples, the simpsec package was used, which employs cluster similarity spectrum CSS.^78^

### Cell-Cell interaction analysis

NicheNet was used to infer interactions between upregulated molecules in the CSF of MS patients and the top 100 upregulated genes in MS versus control ependymal cells.^44^ The Nichenet_output$ligand_activity_target_heatmap function was used to visualize ligand regulatory activity. Ligand-target gene interactions with a weight of less than 0.05 were filtered.

### RNA-Regressing Principal components for the Assembly of Continuous Trajectory (RePACT)

Following Seurat standard pipelines (i.e. normalizing, scaling, and PCA analysis), RePACT was applied to infer genes that vary along disease associated regression trajectories.^37^ The RePACT function plot_scRNA_RePACT_heatmap was used to visualize statistically significant genes (q-value < 0.05) varying along this trajectory, organized into low-to-high binned pseudostates. The distribution of cells within the first three principal components utilized by RePACT was represented using the scatter3D function from the plot3D package in R.

### Motif Enrichment Analysis

First, a position weight matrix indicating TF binding motifs was obtained from the JASPAR 2020 database.^79^ Next, using the AddMotifs function in Signac, the motif information was added to the ATAC assay. Using the RunChromVAR function, motif activity was calculated for each cell and added to a new "chromvar" assay. Finally, the FindAllMarkers function from Seurat was applied to the "chromvar" assay to calculate significant differentially expressed motifs (DEMs) between MS and control cells.

### Inferring Gene Regulatory Networks from Multiomic Data

To identify gene regulatory networks (GRNs) acting in control and MS, the standard SCENIC+ workflow was applied.^41^ Using Pycistopic topic modeling, candidate enhancers and co-accessible regions were identified within their respective conditions. Next, with Pycistarget, DEMs were inferred within differentially accessible regions between control and MS. These two results were combined with gene expression data using a gradient-boosting machine learning approach to uncover differentially active gene regulatory networks. The downstream analysis focused on TF regulons with a positive relationship between region-gene and TF-gene links and a significantly different Eregulon Specific Score (eRSS) between MS and control (p.adj < 0.05).

### Cytoscape Visualization

To visualize the gene regulatory networks constructed by SCENIC+, TF regulons, along with their corresponding region-gene links and associated importance_TF2G values, were extracted from the output MuData object. The extracted data were saved in a CSV file and imported into Cytoscape,^80^ where node colors were mapped to TF regulons and edge transparency was mapped to the importance_TF2G values of the region-gene links.

### Electron microscopy

Periventricular tissue used for EM was extracted at the CHUM from the brains of 2 patients with MS (MS6 and MS7; Supp. Table 3) within 5 hours post-mortem and immediately drop-fixed in a solution of 2.5% glutaraldehyde, 2% paraformaldehyde, and 0.1M sodium cacodylate in double-distilled water (ddH_2_O) for 24-48 hours at 4 °C, before transferring PBS. Tissues were cycled into fresh PBS once per week at 4 °C until processing. Once transported to the Facility for Electron Microscopy Research (FEMR) at McGill University, tissues were transferred back to the glutaraldehyde-based fixative solution with 4% sucrose overnight at 4 °C, and washed 3x by aspiration with 0.1M cacodylate washing buffer. Samples were post-fixed with 1% aqueous osmium tetroxide plus 1.5% aqueous potassium ferrocyanide for 2 hours at 4 °C, followed by three washes with ddH_2_O. Samples were then dehydrated in a standard set of acetone baths of increasing concentration at room temperature (RT) and subsequently immersed in 1:1 Epon:acetone overnight, 2:1 Epon:acetone the next day, and 3:1 Epon:acetone the following night, all on a rotator at RT. Tissues were then infiltrated with 100% Epon for four hours at RT with tube caps off: two hours on a rotator, and 2 hours under a vacuum. Specimens were then embedded in molds, left at RT for 1 hour in a fume hood, and polymerized in an oven at 60 °C for 48 hours. Once polymerized, samples were removed from the oven, cooled at RT, and sectioned. Sections were stained with toluidine blue to confirm tissue integrity. Separate 100nm-thick counter-stained sections were placed on TEM grids for imaging. TEM grids were imaged on a FEI Tecnai G2 Spirit Twin 120 kV Cryo-TEM.

### Mouse Ependymal Cell Culture

Following full gestation and birth, C57BL/6 mouse neonates were collected (typically between 6-8 animals). The neonates (P0-P4) were sacrificed (decapitation) and the deep brain matter (ventricular zone/periventricular zone) was isolated using sterilized sharp-nosed forceps and scalpel blades. Following dissection, the isolated brain tissue was placed in ice-cold sterile PBS (25mL). While in PBS, the tissue was further partitioned via a microsurgical knife into ∼2mm by 2mm sized pieces to increase surface area for enzymatic digestion. Tissue was extracted from PBS via passage through a 40 µm mesh strainer and digested in 20-25mL of enzymatic solution (TrypLE) in a 37°C water bath. After 15-20 minutes, the tissue was removed from the water bath and mechanically triturated 8-10 times via pipette. The tissue was returned to the water bath for a further 15-20 minutes, removed, and triturated again, and placed back in the water bath. After 5-10 more minutes, the enzymatic reaction was quenched using an equal volume of media containing 15% fetal bovine serum (FBS), 2% penicillin-streptomycin (P/S) and 1% Glutamax. The mixture was fed through a 40 µm mesh strainer and into a 50ml tube then centrifuged at 350g for 10 minutes. Following centrifugation, supernatant was discarded, and cells were counted via hemocytometer. Cells were resuspended in media containing 10% FBS, 1% P/S and 1% Glutamax (“proliferation media”) to a target density of 500-600,000 cells/mL. They were then plated onto poly-d-lysine coated cell culture vessels (96-well optic bottom plates for use in immunocytochemistry) and left to rest for 3 days. Media was changed to proliferation media at day 3 to remove debris, then after a further 2-4 days (5-7 days total in proliferative conditions) cells were deemed confluent. Confluent cells were then placed in media without FBS (1% P/S, 1% Glutamax, and 0.25 µM small molecule MLN-4924 (“differentiation media 1”; DM1). Media was changed after 3 days with DM1. After 6 days in DM1, media was changed to include 0.5% FBS, 1% P/S, 1% Glutamax and 0.125 µM small molecule MLN-4924 (“differentiation media 2”; DM2) to encourage a mature ependymal phenotype. Cells were left for nine days in DM2 (with a media change every 3 days). By this stage, cultures display mature ependymal cell markers – approximately 60-70% of the cells are FOXJ1 positive ependymal cells. Cytokines were added for 4 days, then cells were post-fixed in 4% PFA for 10 minutes, followed by immunocytochemistry. After washes with 1x PBS, cells were blocked with 10% donkey serum and 0.3% Triton-X at room temperature. After blocking, cells were incubated with primary antibodies (FOXJ1; 1:250, Fisher Scientific, 501128960) diluted in 1x PBS with 4% donkey serum and 0.3% Triton-X overnight at 4°C. After washes with PBS with 0.3% Triton-X, cells were incubated with Alexa Fluor secondary antibodies diluted in 1x PBS. After washes with 1x PBS, nuclei were stained with DAPI (1:5000; cat# D1306; Invitrogen). The Molecular Devices MetaXpress high-content screening platform was used for imaging. Eight images per well were captured at 20x magnification and analyzed. Image locations were kept consistent across plates. Analysis parameters were optimized to exclude outliers and debris. FOXJ1 staining was co-localized with the DAPI-positive nuclei. Cells with staining intensity above background were considered positive.

### Animals

Ifngr1^fl/fl^-Cre^ERT2^-lsl-Tdt animals were sourced from the laboratory of Dr. Alexander Gregorieff and genotyped according to standard PCR and gel electrophoresis protocols (**Supp. Figure 6A**). While Clu shows low levels of expression in all tissues, it is primarily expressed in epithelial cells (including ependymal cells and choroid plexus epithelial cells), allowing its promoter to be a useful driver for targeted ependymal effects,^81^ especially in an acute context. This has been further supported by the fact that our own scRNAseq data has confirmed that it is highly expressed in ependymal cells.^33^

### Intracerebroventricular (ICV) injections

ICV injections were conducted as previously described.^33^ Briefly, induction anesthesia parameters were 5% isoflurane (1.5 L/min oxygen) and initial maintenance parameters were 2.5% isoflurane (0.8L/min oxygen). Injection coordinates to target the lateral ventricle were the same as previously described (AP −0.25mm, LM 0.85mm, DV −2.5mm) and injection volume was 2uL. For insult injections, the rate was 0.25uL/min and the needle was held in place for 5 minutes prior to injection, for 2 minutes at −1.25mm DV during needle removal, and 7 minutes post-injection. For dextran injections, the rate was 0.067uL/min and the needle was held in place for 5 minutes prior to injection and 15 minutes post-injection. For non-recoverable injections, euthanyl was administered at a dosage 0.05mL per 10g of body weight at a concentration of 24mg/mL post-injection.

## Acknowledgements

The authors wish to thank the staff at the <u>McConnell Brain Imaging Centre</u> (BIC) at The Neuro for their support of our *ex vivo* human brain slab scan optimization, Hong Li for her expertise in human brain tissue processing and sectioning, and Kelly Sears and Johanne Ouellette of the <u>Facility for Electron Microscopy Research</u> (FEMR) at McGill for processing human tissue for electron microscopy. They also extend gratitude to Dr. Alexander Gregorieff for providing the Ifngr1^fl/fl^-Cre^ERT2^-lsl-Tdt animals, to the staff at the <u>Center for Neurological Disease Models</u> (CNDM) at The Neuro for taking care of animals prior to surgery, and to Vanessa Omana for her contribution to extraction of single nuclei for sequencing. They are also grateful for feedback provided from Drs. G.R. Wayne Moore and Jack P. Antel. Lastly, they offer sincere appreciation to the <u>Douglas Bell Canada Brain Bank</u> (DBCBB), and to the donors themselves for their valued contribution to science. They made the majority of this work possible.

