## Supplemental Figures for "An MRI-informed histo-molecular analysis implicates ependymal cells in the pathogenesis of periventricular pathology in multiple sclerosis"

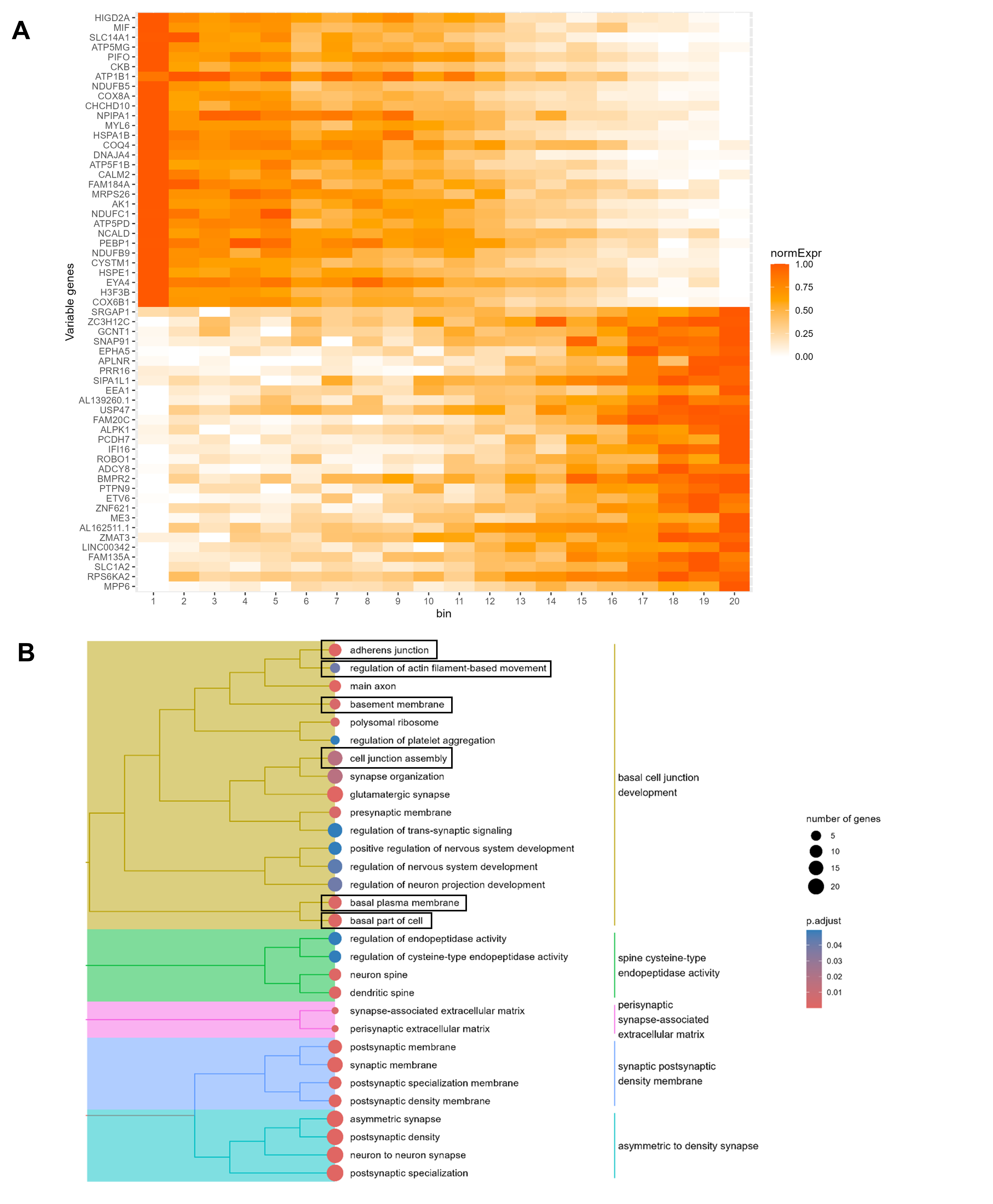

**Supp. Figure 1**

**Supplemental Figure 1. (A)** Heatmap showing normalized expression of the top 30 statistically significant genes (q-value < 0.05) per condition (control – top 15; MS – bottom 15) along a logistic regression-PCA-derived MS trajectory, organized into low-to-high binned pseudostates. Bin 1 represents the least diseased cells (control), while bin 20 represents the most diseased cells (MS). Colour represents normalized gene expression (0-1). **(B)** Gene ontology (GO) enrichment plot illustrating GO terms upregulated in MS-associated ependymal cells (associated with basic differential gene expression testing conducted in Seurat). Note terms associated with cell junction assembly (black boxes), similar to the gene ontology of enriched terms in MS generated by differential gene expression in Figure 3F. Dot size represents the number of genes associated with a GO term, whereas dot colour represents adjusted p-value (p.adjust).

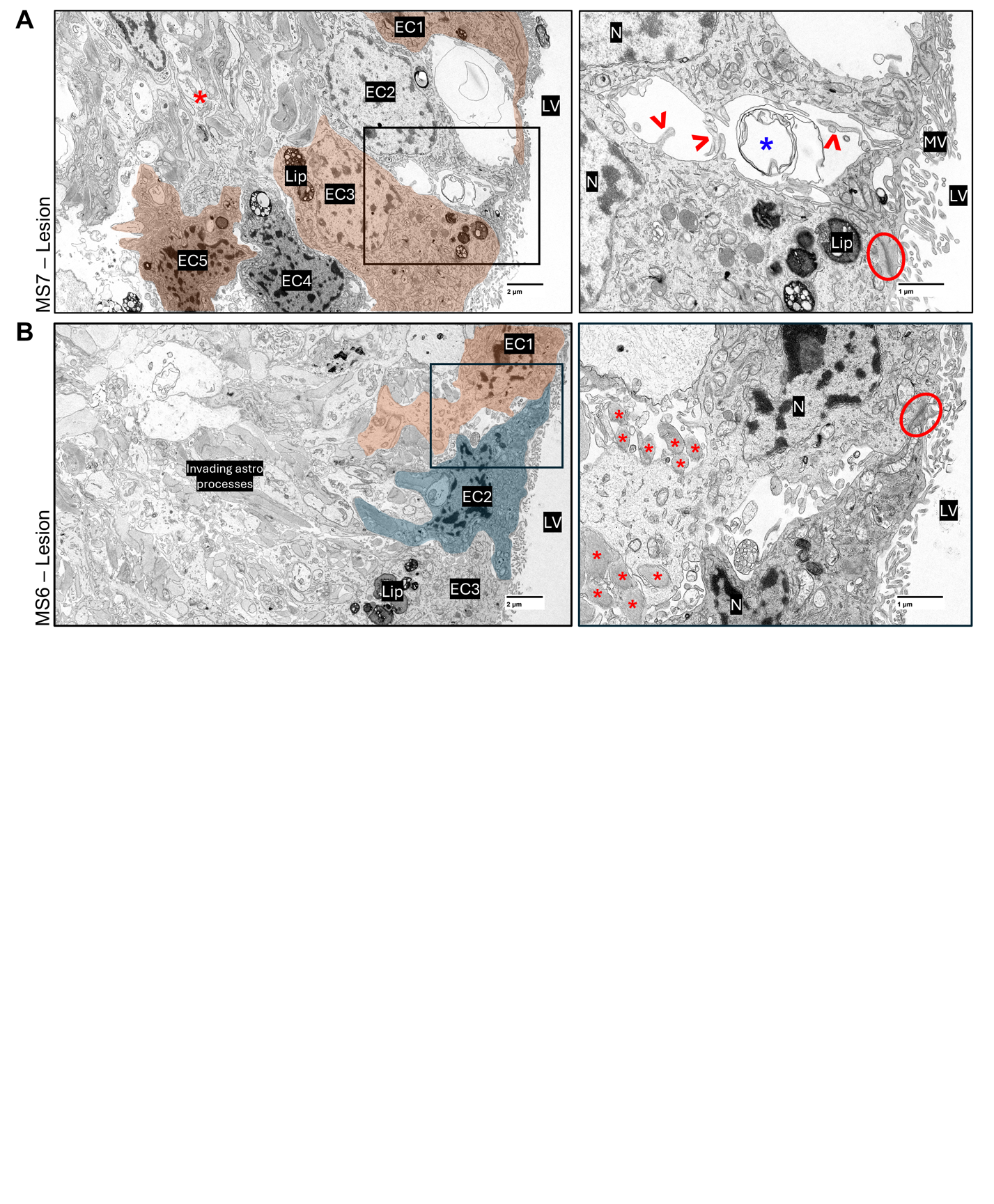

**Supplemental Figure 2. (A)** (Left) Image of the ependymal lining (MS7), adjacent to a periventricular lesion, demonstrating ependymal cell adhesion disruption. A black box indicates the region of the magnified panel on the right. A red star indicates subependymal astroglial processes. Scale bar = 2µM. (Right) Magnified image of lesion-proximal ependymal cells with disrupted cell-cell adhesion. Red arrowheads point to lateral cellular processes ‘reaching out’ from detached ependymal cells and a blue star denotes a circular structure present in the space between separated ependymal cells. A red circles denotes an adherens junction. Scale bar = 1µM.

**(B)** (Left) Image of the ependymal lining in an MS brain (MS7), proximal to a periventricular lesion, demonstrating highly irregular cell morphology (EC1 – orange-highlighted; EC2 – blue-highlighted). Also note the considerable lipofuscin buildup in EC3. A black box indicates the region of the magnified panel on the right. Scale bar = 2µM. (Right) Magnified image of lesion-proximal ependymal cells with irregular morphology. A red circle denotes an adherens junction, and red stars indicate astrocyte processes that have invaded the intercellular spaces formed by ependymal cell-cell adhesion disruption, suggesting barrier function compensation. Scale bar = 1µM. EC = ependymal cell; N = nucleus; MV = microvilli; LV = lateral ventricle; Lip = lipofuscin.

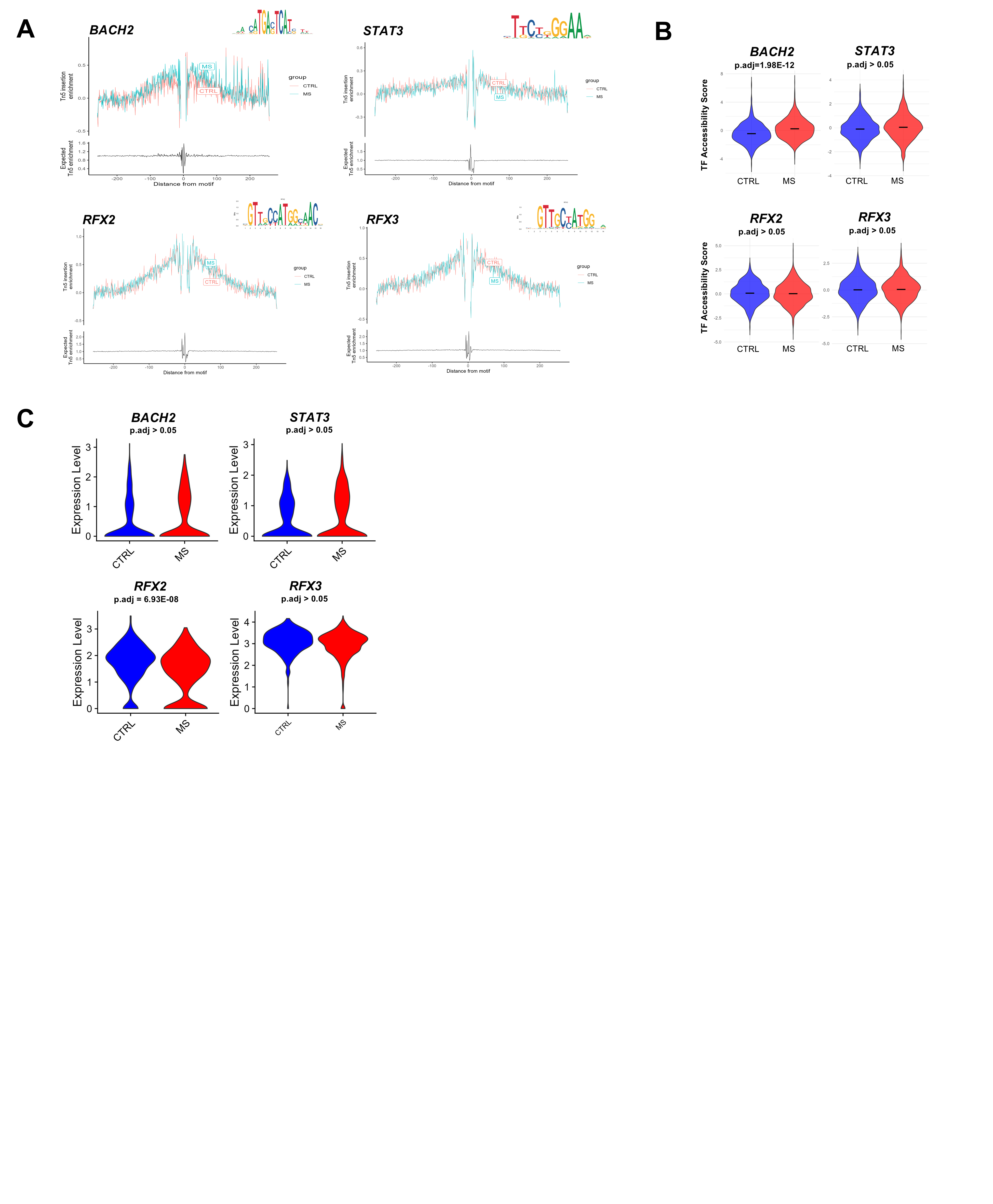

**Supplemental Figure 3. (A)** DNA footprinting analysis plots for the TFs with the top scaled eRSS values in control and MS conditions: *BACH2*, *STAT3*, *RFX2*, and *RFX3*. **(B)** Motif accessibility plots *BACH2*, *STAT3*, *RFX2*, and *RFX3*. **(C)** Gene expression violins for *BACH2*, *STAT3*, *RFX2*, and *RFX3*.

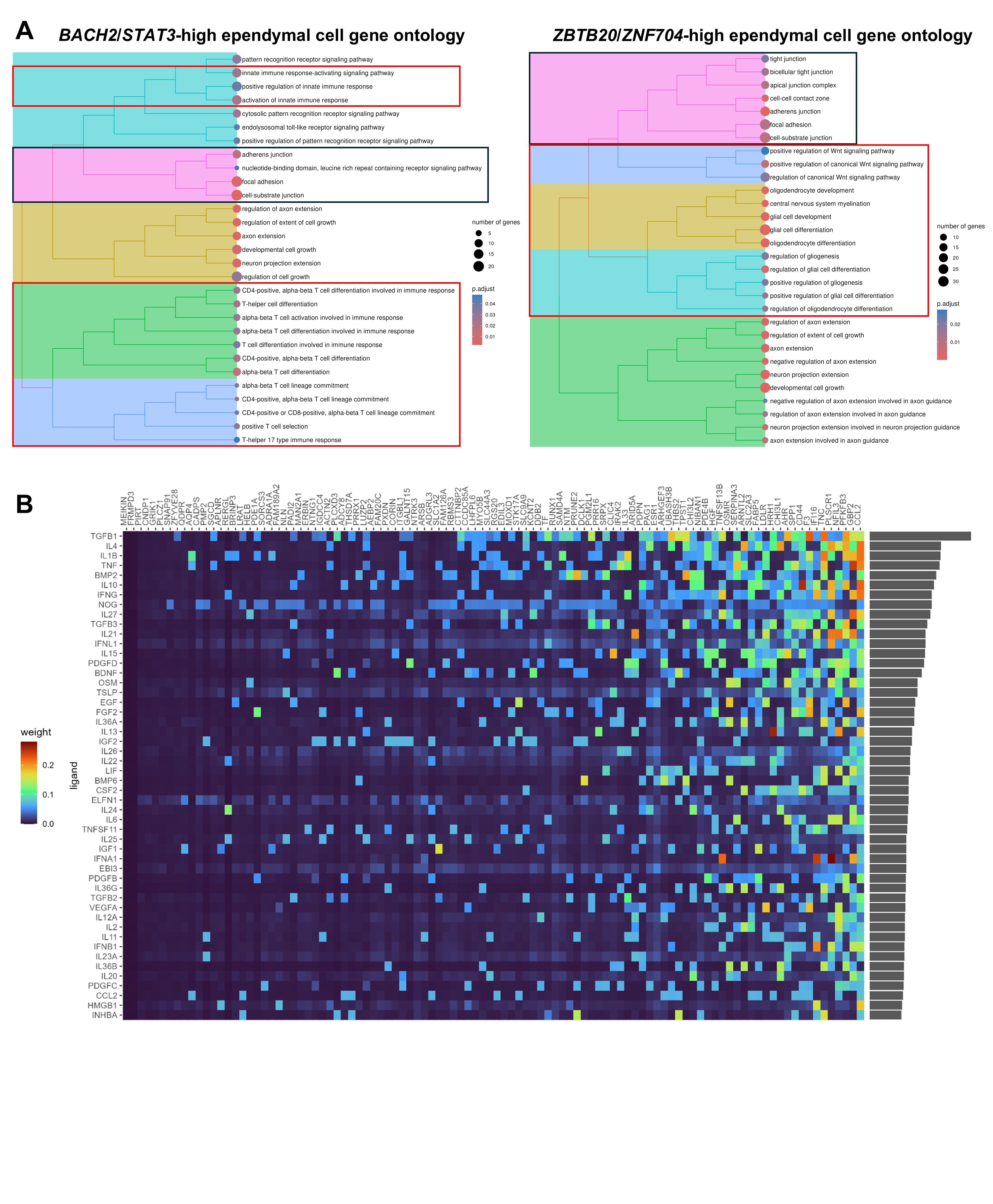

**Supp. Figure 4**

**Supplemental Figure 4. (A)** (Left) Gene ontology analysis of MS ependymal cells enriched in *BACH2*/*STAT3* expression compared to control ependymal cells demonstrated upregulation of unique pathways related to immune regulation and inflammation compared to MS ependymal cells enriched in *ZBTB20*/*ZNF704* activity (red boxes). (Right) Conversely, gene ontology analysis of MS ependymal cells enriched in *ZBTB20*/*ZNF704* expression compared to control ependymal cells demonstrated upregulation of unique pathways related to WNT signaling, glial cell differentiation, and myelination compared to ependymal cells enriched in *BACH2*/*STAT3* expression. Both populations of ependymal cells demonstrated upregulation of pathways related to cell junction assembly and cell morphology and/or cytoplasmic alterations (black boxes). Dot size represents the number of genes associated with a GO term, whereas dot colour represents adjusted p-value (p.adjust). **(B)** Heatmap of all ligands that were predicted to drive upregulated genes in MS ependymal cells. Colour indicates predicted interaction weight.

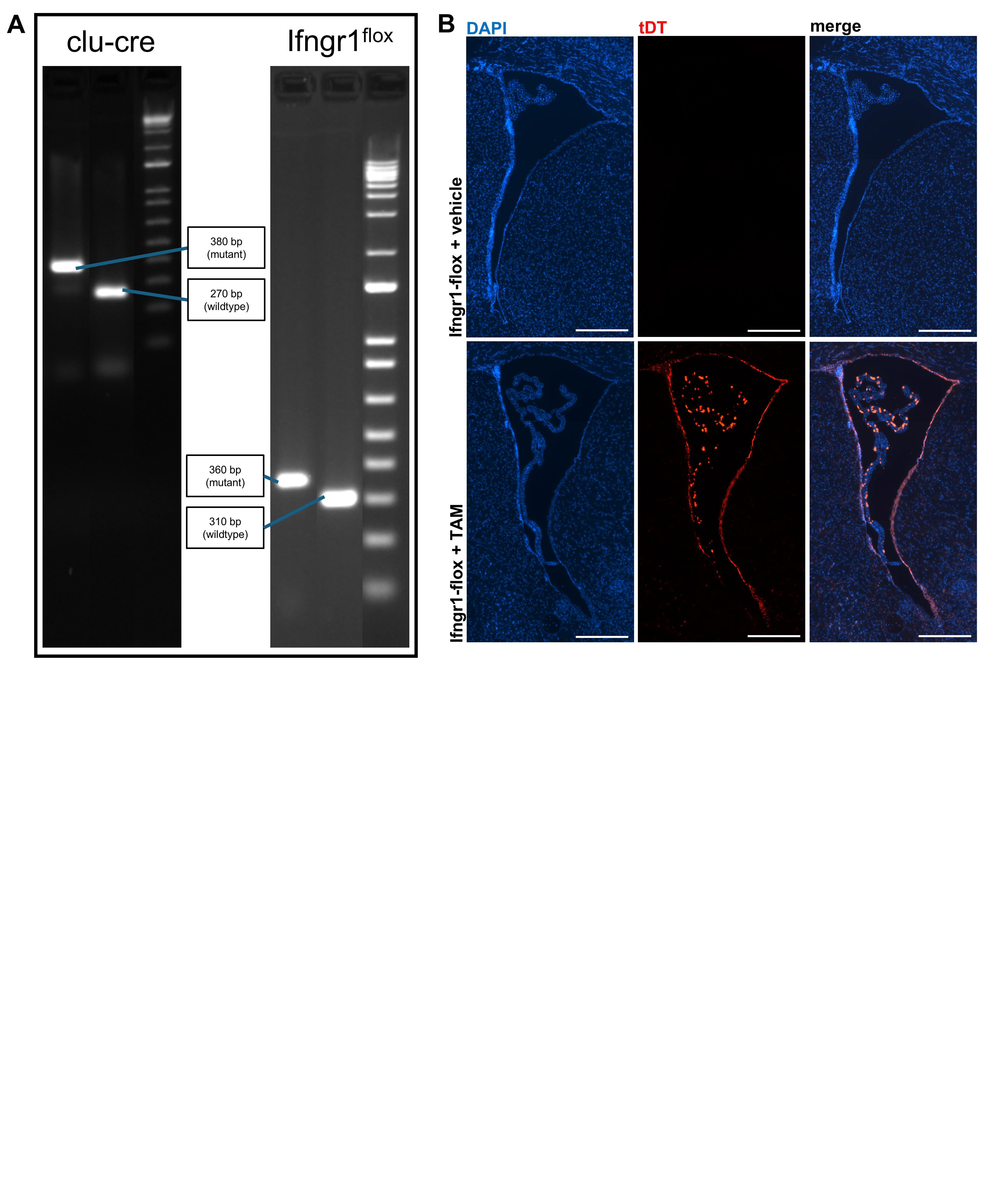

**Supplemental Figure 5. (A)** Representative images of gels obtained from genotyping clu-cre/lsl-tdt and Ifngr1-floxed animals, showing relevant clu-cre and IFNGR1 bands, respectively. **(B)** Stitched fluorescent images of the right lateral ventricle in vehicle-administered versus tamoxifen (TAM)-administered Ifngr1^fl/fl^-Cre^ERT2^-lsl-Tdt animals. Vehicle-administered animals showed no cre recombination/TdTomato (TdT) expression, whereas TAM-administered animals showed clear cre recombination/Tdt expression in ependymal cells and choroid plexus epithelial cells. Scale bars = 100µM.
