## Supplemental Table 1 for "An MRI-informed histo-molecular analysis implicates ependymal cells in the pathogenesis of periventricular pathology in multiple sclerosis"

|  |  |  | Angular bands contained in histological tissue block(s) | |
| --- | --- | --- | --- | --- |
| Brain ID | Slab ID | Position of Slab in Brain | Ependyma-in gradient | No gradient |
| MS #1 | MS 1F | Frontal | N/A | 2-4 |
|  | MS 1M | Middle | N/A | 3, 4 |
|  | MS 1C | Caudal | N/A | 2, 3 |
| MS #2 | MS 2F | Frontal | 2-5 (Block #1)  11, 12 (Block #2) | 14-16 |
|  | MS 2M | Middle | N/A | 4-6 |
|  | MS 2C | Caudal | 2-5 | 23, 24 |
| MS #3 | MS 3F | Frontal | 3, 4 (Block #1)  9-12 (Block #2) | 23, 24 |
|  | MS 3M | Middle | 7-10 | 1, 23, 24 |
|  | MS 3C | Caudal | 4-6 | 11, 12 |
| MS #4 | MS 4F | Frontal | 1-6 (Block #1)  7-13 (Block #2) | 22, 23 |
|  | MS 4M | Middle | N/A | 1-8 |
|  | MS 4C | Caudal | 1-3 (Block #1)  9-11 (Block #2) | 13-15 |
| HC #1 | HC 1F | Frontal | N/A | 1-4 |
|  | HC 1M | Middle | N/A | 4-11 |
|  | HC 1C | Caudal | N/A | 1-10 |
| HC #2 | HC 2F | Frontal | N/A | 5-14 |
|  | HC 2M | Middle | N/A | 2-11 |
|  | HC 2C | Caudal | N/A | 2-8 |
| HC #3 | HC 3F | Frontal | 2-13 | 15-24 |
|  | HC 3M | Middle | N/A | 3-11 |
|  | HC 3C | Caudal | N/A | 17-23 |
| HC #4 | HC 4F | Frontal | 1-5 | 9-14 |
|  | HC 4C | Caudal | N/A | 5-10 |

**Supplemental Table 1**: Brain slab ID information and list of angular bands covered in resected tissue blocks for histology that exhibit an ependyma-in gradient of T1 and T2* relaxation times in periventricular white matter or no gradient. The abbreviation MS refers to the brain of an individual with multiple sclerosis and the abbreviation HC refers to the brain of a healthy individual with no inflammatory or neurodegenerative disease.
