## Supplemental Table 2 for "An MRI-informed histo-molecular analysis implicates ependymal cells in the pathogenesis of periventricular pathology in multiple sclerosis"

| Slab ID | Angular band number | Total change in T_1_, ΔT_1_ (ms) | Total change in T_2_^*^, ΔT_2_^*^ (ms) |
| --- | --- | --- | --- |
| MS 1F | 5 | -29.7 | -5.9 |
| MS 1M | N/A | N/A | N/A |
| MS 1C | 11 | -21.1 | 1.0 |
|  | 12 | -31.2 | -0.4 |
| MS 2F | 2 | -41.7 | -5.8 |
|  | 3 | -93.3 | -15.0 |
|  | 4 | -87.0 | -14.4 |
|  | 5 | -54.4 | -11.7 |
|  | 11 | -52.5 | -8.4 |
|  | 12 | -69.0 | -11.8 |
|  | 20 | -25.9 | -5.5 |
| MS 2M | 6 | -39.3 | -5.2 |
|  | 7 | -21.7 | -3.1 |
|  | 9 | -37.7 | -5.6 |
| MS 2C | 1 | -20.8 | -3.2 |
|  | 2 | -42.8 | -5.1 |
|  | 3 | -51.4 | -6.8 |
|  | 4 | -27.0 | -1.7 |
|  | 5 | -11.8 | -0.9 |
|  | 6 | -47.0 | -3.2 |
|  | 7 | -65.8 | -7.3 |
|  | 8 | -38.2 | -2.8 |
|  | 9 | -34.0 | -3.0 |
|  | 10 | -24.0 | -0.8 |
|  | 11 | -34.2 | -3.2 |
|  | 12 | -30.7 | -3.3 |
| MS 3F | 3 | -31.1 | -4.1 |
|  | 4 | -43.1 | -7.3 |
|  | 11 | -45.3 | -4.8 |
|  | 12 | -20.4 | -0.7 |
| MS 3M | 7 | -27.7 | -4.2 |
|  | 8 | -65.8 | -6.4 |
|  | 9 | -81.5 | -7.0 |
|  | 10 | -94.0 | -10.6 |
|  | 11 | -47.5 | -5.4 |
| MS 3C | 4 | -16.7 | -1.4 |
|  | 5 | -32.8 | -2.4 |
|  | 6 | -13.3 | -1.5 |
| MS 4F | 1 | -77.2 | -8.9 |
|  | 2 | -94.4 | -9.9 |
|  | 3 | -87.7 | -9.1 |
|  | 4 | -94.0 | -9.0 |
|  | 5 | -99.7 | -8.1 |
|  | 6 | -109.6 | -7.1 |
|  | 7 | -101.5 | -6.2 |
|  | 8 | -72.0 | -3.5 |
|  | 9 | -81.0 | -3.2 |
|  | 10 | -86.9 | -3.9 |
|  | 11 | -100.3 | -6.6 |
|  | 12 | -120.0 | -12.1 |
|  | 13 | -119.1 | -18.0 |
|  | 24 | -75.1 | -10.6 |
| MS 4M | 8 | -40.1 | -5.1 |
|  | 9 | -32.8 | -4.5 |
| MS 4C | 1 | -84.8 | -12.4 |
|  | 2 | -116.6 | -14.1 |
|  | 3 | -135.1 | -15.7 |
|  | 5 | -113.8 | -10.1 |
|  | 8 | -42.9 | -3.6 |
|  | 9 | -29.5 | -2.3 |
|  | 10 | -92.5 | -9.0 |
|  | 11 | -65.3 | -6.4 |
| HC 1F | 1 | -21.5 | -3.4 |
|  | 4 | -30.6 | -4.6 |
|  | 6 | -32.0 | -3.6 |
|  | 8 | -26.5 | -2.92 |
|  | 9 | -23.6 | -3.8 |
|  | 24 | -26.5 | -2.9 |
| HC 1M | 7 | -5.3 | -3.4 |
|  | 10 | -44.4 | -6.6 |
|  | 11 | -45.1 | -6.3 |
| HC 1C | 1 | -27.0 | -5.4 |
|  | 2 | -34.5 | -5.7 |
|  | 3 | -30.2 | -5.6 |
|  | 4 | -26.3 | -3.5 |
|  | 7 | -7.0 | -2.4 |
|  | 8 | -7.0 | -2.2 |
|  | 10 | -14.7 | -3.4 |
|  | 11 | -8.9 | -2.3 |
| HC 2F | N/A | N/A | N/A |
| HC 2M | 2 | -13.2 | -3.1 |
|  | 3 | -19.1 | -3.1 |
|  | 5 | -24.7 | -5.3 |
|  | 6 | -22.4 | -5.5 |
| HC 2C | 2 | -22.9 | -2.6 |
|  | 3 | -24.9 | -3.6 |
|  | 4 | -31.9 | -4.0 |
|  | 5 | -12.9 | -4.9 |
|  | 6 | -10.1 | -3.7 |
|  | 7 | -18.2 | -4.0 |
|  | 8 | -0.6 | -2.5 |
|  | 9 | -27.5 | -3.7 |
|  | 10 | -27.6 | -4.0 |
|  | 11 | -23.5 | -3.8 |
|  | 13 | -21.5 | -2.8 |
| HC 3F | 2 | -15.2 | 0.6 |
|  | 3 | -54.7 | -1.7 |
|  | 4 | -41.6 | -1.9 |
|  | 5 | -43.8 | -0.6 |
|  | 6 | -38.4 | -0.1 |
|  | 7 | -41.6 | -1.4 |
|  | 8 | -45.7 | -2.9 |
|  | 9 | -54.9 | -4.4 |
|  | 10 | -47.9 | -2.7 |
|  | 13 | -43.4 | -5.3 |
| HC 3M | N/A | N/A | N/A |
| HC 3C | 1 | -11.0 | -2.2 |
|  | 2 | -26.5 | -3.9 |
|  | 6 | -21.6 | -0.6 |
|  | 13 | -13.2 | -1.6 |
|  | 23 | -9.7 | -2.9 |
|  | 24 | -27.4 | -4.9 |
| HC 4F | 1 | -4.6 | -1.3 |
|  | 2 | -13.2 | 0.8 |
|  | 3 | -34.6 | -1.7 |
|  | 4 | -22.1 | -2.9 |
|  | 12 | -43.3 | -9.2 |
| HC 4C | 1 | -24.3 | -3.8 |
|  | 3 | -18.3 | -2.1 |
|  | 4 | -16.2 | -1.4 |
|  | 9 | -17.3 | -1.4 |
|  | 10 | -17.2 | -1.0 |

**Supplemental Table 2**: Total change in T1 or T2* over ependyma-in reductions in relaxation time observed in angular bands of MS or control brain slabs. Some angular bands were omitted, despite showing ependyma-in reductions in relaxation times, owing to non-negligible partial volume effects with bordering cortical gray matter.
