## Supplemental Table 3 for "An MRI-informed histo-molecular analysis implicates ependymal cells in the pathogenesis of periventricular pathology in multiple sclerosis"

| Brain ID | Implicated  Experiments | Sex | Age | Disease Duration | MS Type | COD | PMD | Tissue Type |
| --- | --- | --- | --- | --- | --- | --- | --- | --- |
| HC1  *DH1143* | MRI  IHC  snRNA-seq | M | 64 | N/A | N/A | Caecal  carcinoma | 20.75h | Formalin-fixed  Fresh-frozen |
| HC2  *DH1670* | MRI  IHC  snRNA-seq | F | 51 | N/A | N/A | Breast cancer; liver metastasis | 26.25h | Formalin-fixed  Fresh-frozen |
| HC3  *DH1722* | MRI  IHC  snRNA-seq | M | 59 | N/A | N/A | Stomach cancer; obstructive hypertrophic cardiomyopathy | 17.67h | Formalin-fixed Fresh-frozen |
| HC4  *DH989* | MRI  IHC  snRNA-seq | M | 79 | N/A | N/A | Cardiac insufficiency; pulmonary edema | 14.75h | Formalin-fixed  Fresh-frozen |
| HC5  *DH1888* | snRNA-seq | F | 72 | N/A | N/A | Cardiomegaly; pulmonary edema; reactive myeloid hyperplasia | 37.42h | Fresh-frozen |
| MS1  *DH1011* | MRI  IHC  snRNA-seq | M | 64 | 31 years | N/A | Pneumonia; carcinoma of vocal chord | 63h | Formalin-fixed  Fresh-frozen |
| MS2  *DH1002* | MRI  IHC  snRNA-seq | M | 77 | 41 years | N/A | Pneumonia | 11.25h | Formalin-fixed  Fresh-frozen |
| MS3  *DH1090* | MRI  IHC  snRNA-seq | M | 73 | 40 years | N/A | Advanced MS; bladder tumor, obstruction of urinary tract | 57.75h | Formalin-fixed  Fresh-frozen |
| MS4  *DH1736* | MRI  IHC  snRNA-seq | F | 58 | 29 years | N/A | Hepatic insufficiency; perforated ulcer, sepsis | 14.05h | Formalin-fixed  Fresh-frozen |
| MS5  *DH1421* | snRNA-seq | F | 46 | 6 years | N/A | Lung cancer | 22.5h | Fresh-frozen |
| MS6  *AB276* | EM | M | 60 | 38 years | SP | MAID | 4.13h | Fresh |
| MS7  *AB280* | EM | M | 56 | 34 years | PP | MAID | 5.25h | Fresh |

**Supplemental Table 3**. Patient Information.
